# Structure–activity–guided design of potent and selective SSTR2 agonists inspired by a cone snail venom peptide

**DOI:** 10.64898/2026.11.24.768804

**Authors:** Ho Yan Yeung, Nicholas Schumann, Alina Anhnhi Vo, Drake Hampton, Colton Gschwandtner, Seyed Reza Moosavi, Paul Schlitterlau, Martin Ullrich, Robert Wodtke, Michael J. Robertson, Helena Safavi-Hemami

## Abstract

The human somatostatin 2 receptor (SSTR2) is an important diagnostic and therapeutic target in neuroendocrine disorders and cancer. Because the endogenous peptide hormone somatostatin lacks *in vivo* stability and receptor subtype selectivity, significant effort has gone into the development of stable and selective peptide ligands such as octreotide and vapreotide. We recently identified Consomatin pG1, a somatostatin mimic from the venomous fish-hunting cone snail *Conus geographus*, that shares key stability-enhancing features with clinical somatostatin analogs, including a compact cyclic fold and a D-Trp residue, while exhibiting superior selectivity for SSTR2. Here, we define the structural basis of this selectivity by determining the cryo-electron microscopy (cryo-EM) structure of the pG1-human SSTR2-Gαi3 complex at 1.9 Å resolution. Combined with receptor chimera and mutagenesis studies, these data reveal how pG1 engages the orthosteric binding site and forms critical interactions with extracellular loop (ECL) regions, notably ECL2. Guided by these insights, we designed a series of pG1 analogs with further enhanced potency, while retaining selectivity at SSTR2. Together, our findings offer insight into the molecular determinants of SSTR2 engagement and selectivity and provide new SSTR2-selective tool compounds and potential therapeutic leads.

## Introduction

Somatostatin is a peptide hormone that exists in two bioactive isoforms, SS-14 and SS-28 (1–3). In humans, it acts as the endogenous ligand for five somatostatin receptor subtypes (SSTR1–5), a family of Gα_i/o_-coupled class A G protein-coupled receptors (GPCRs) expressed across diverse tissues (1, 4). Through activation of distinct receptor subtypes, somatostatin regulates a wide range of physiological processes. For example, somatostatin controls glucose homeostasis by suppressing glucagon and insulin secretion via SSTR2 and SSTR5 activation in pancreatic islets (5–8). In the nervous and immune systems, somatostatin signals through SSTR1 and SSTR4 to modulate nociception and immune responses (9–12).

Because somatostatin displays poor metabolic stability and lacks receptor selectivity, substantial efforts have focused on developing stable somatostatin analogs with improved pharmacological profiles (13, 14). Several cyclic peptide analogs, such as octreotide and vapreotide, have achieved clinical success, particularly through targeting SSTR2 (15–17). These analogs are widely used for the treatment of neuroendocrine disorders such as acromegaly (18) and have also been applied in diagnostic imaging and radionuclide therapy in oncology (15–17). Despite these advances, currently available SSTR2-directed ligands, while effective, retain residual activity at other receptor subtypes, providing opportunities for further improvement. For example, compared to somatostatin, octreotide shows enhanced stability and preferential activity at SSTR2 but retains measurable activity at other receptor subtypes, most notably SSTR3 and SSTR5 (12). This residual cross-reactivity reflects the high sequence and structural conservation among SSTR subtypes and may represent a limitation for both therapeutic precision and mechanistic understanding.

We previously identified the somatostatin-like toxin Consomatin nG1 from the fish-hunting cone snail *Conus geographus*, from which the bioactive peptide Consomatin pG1 (hereafter pG1) is derived, and showed that pG1 potently suppresses pancreatic glucagon secretion via SSTR2 activation (19). Consomatin pG1 shares striking similarity with clinically used somatostatin analogs, particularly octreotide, including the presence of a D-Trp and a compact, stability-enhancing cyclic fold (**Fig. 1A**). However, unlike octreotide, pG1 exhibits improved SSTR2 selectivity (19). These properties make pG1 a valuable molecular probe for elucidating the structural basis of SSTR subtype selectivity and for guiding the design of next-generation SSTR2-selective ligands.

**Figure 1:**
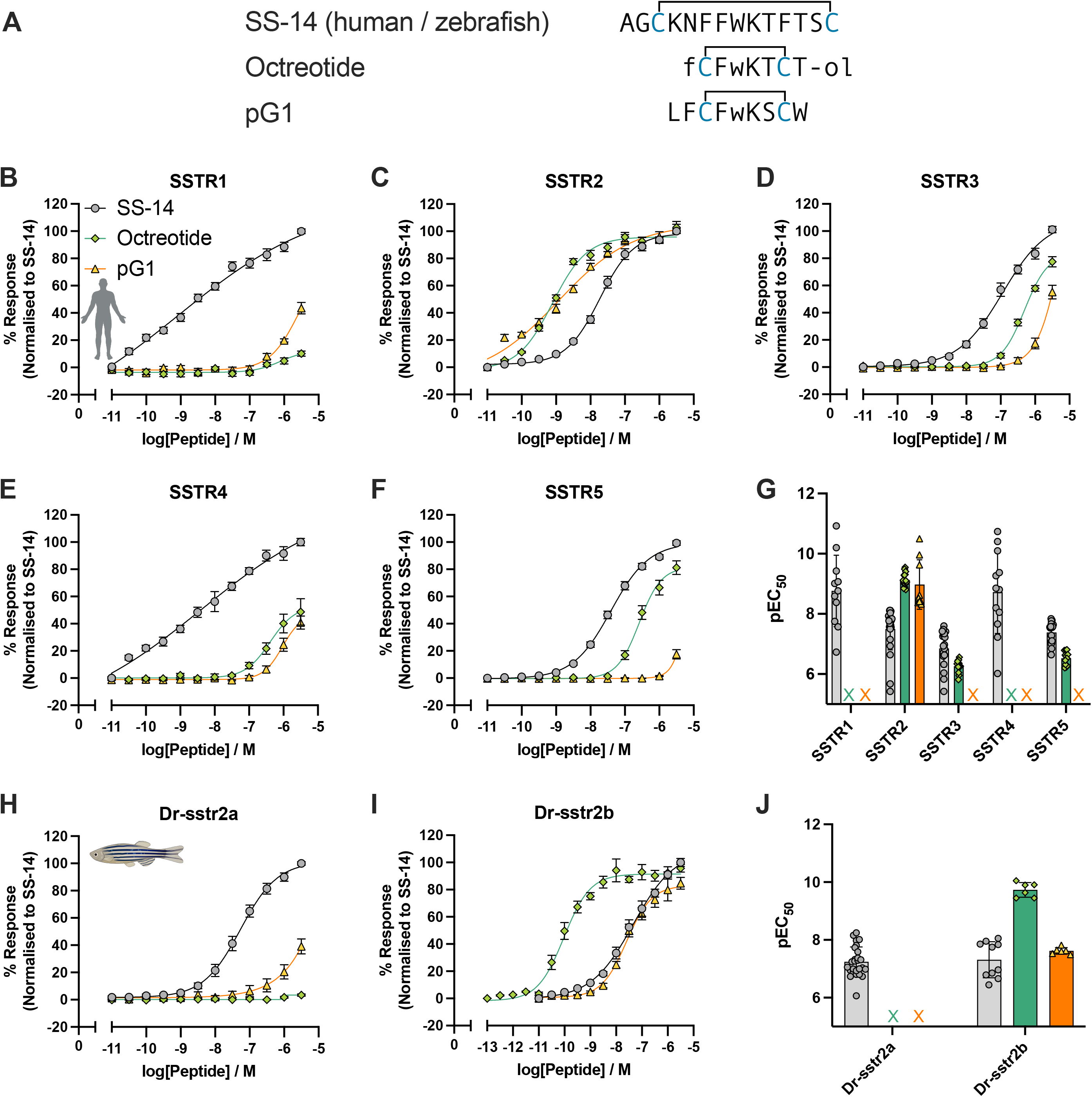
pG1 exhibits enhanced selectivity for human SSTR2 compared with octreotide. (A) Sequence alignment of SS-14, octreotide and pG1. f, d-phenylalanine; w, d-tryptophan, -ol, alcohol group. Disulfide bonds are indicated with bracket, and the cysteines highlighted in blue. (B-F) Representative concentration–response curves illustrating activation of human SSTR1–5 by SS-14, octreotide, and pG1 using the PRESTO-Tango β-arrestin recruitment assay. All data are presented as mean ± S.E.M. from three independent biological replicates, each performed with technical duplicates. (G) Scatter plot summarizing pEC50 values of SS-14, octreotide and pG1 at hSSTR1-5 (mean ± S.D.) (H-I) Representative concentration–response curves showing activation of zebrafish Dr-sstr2a and Dr-sstr2b by octreotide and pG1. (J) Scatter plot summarizing pEC_50_ values for SS-14, octreotide, and pG1 at Dr-sstr2a and Dr-sstr2b (mean ± S.D.). X indicates weak (< 50% Emax) or no receptor activation.

Notably, we previously also observed that pG1 potently activates the zebrafish SSTR2 homolog Dr-sstr2b but shows little to no detectable activity at the closely related isoform Dr-sstr2a (19, 20). This finding was surprising given the strong conservation of the orthosteric ligand-binding pocket among human SSTR2, Dr-sstr2a, and Dr-sstr2b. The lack of activity at Dr-sstr2a suggests that determinants outside the orthosteric binding site are critical for receptor activation and species-specific selectivity. Importantly, this divergence provides an opportunity to leverage the differential responsiveness of zebrafish receptors to investigate the contribution of non-orthosteric regions to ligand engagement using chimeric receptor mutants.

Here, to define the structural features underlying pG1’s SSTR2 selectivity, we combined targeted receptor mutagenesis, including extracellular loop (ECL)-swaps and point mutations with high-resolution cryo-EM structure of pG1 bound to human SSTR2 in complex with Gαi3. These integrated approaches reveal that pG1 engages human SSTR2 through a network of interactions extending beyond the transmembrane core, involving ECL regions, particularly ECL2 and ECL3. Differences within these extracellular regions explain the divergent pharmacological responses of human and zebrafish SSTR2 receptors and support the previously described role of extracellular regions in SSTR activation. Together, these findings provide a structural framework for understanding species- and subtype-specific somatostatin receptor engagement and for guiding the design of next-generation SSTR2-selective ligands with improved potency and selectivity.

## Results

### pG1 potently and selectively activates human SSTR2 and zebrafish Dr-sstr2b but has almost no detectable activity at zebrafish Dr-sstr2a

To assess human SSTR subtype and zebrafish Dr-sstr2 selectivity, we employed the PRESTO-Tango β-arrestin recruitment platform (21). As previously reported, human SS-14 activated all 5 human SSTR subtypes, and both octreotide and pG1 acted as potent full agonists at SSTR2, with EC_50_ values of 18.5 nM (pEC_50_ ± S.E.M.: 7.73 ± 0.04) for SS-14, 0.78 nM (pEC_50_ ± S.E.M.: 9.11 ± 0.05) for octreotide, and 1.14 nM (pEC_50_ ± S.E.M.: 8.95 ± 0.19) for pG1 (**Fig. 1B-G, Table S1**). In contrast to octreotide, which retained measurable activity at SSTR3 and SSTR5, pG1 showed no detectable activation of SSTR5, and only minimal activity at SSTR3 at concentrations exceeding 1 μM (**Fig. 1B-G**). Furthermore, as previously reported (19), pG1 potently activated the zebrafish SSTR2 homolog Dr-sstr2b but elicited little to no detectable activation of the closely related isoform Dr-sstr2a, with similar results observed for octreotide (**Fig. 1H-J**). Despite high overall similarity and conservation of orthosteric transmembrane binding-site residues between human SSTR2 and Dr-sstr2a, the two receptors differ notably in their ECL sequences (**Fig. 2A, Fig. S1**). This combination of conserved orthosteric architecture and divergent extracellular regions makes Dr-sstr2a a useful comparative template for dissecting the contribution of ECLs to ligand responsiveness. Accordingly, we generated ECL-swap chimeras between human SSTR2 and Dr-sstr2a to directly assess the role of ECL in pG1 responsiveness.

**Figure 2:**
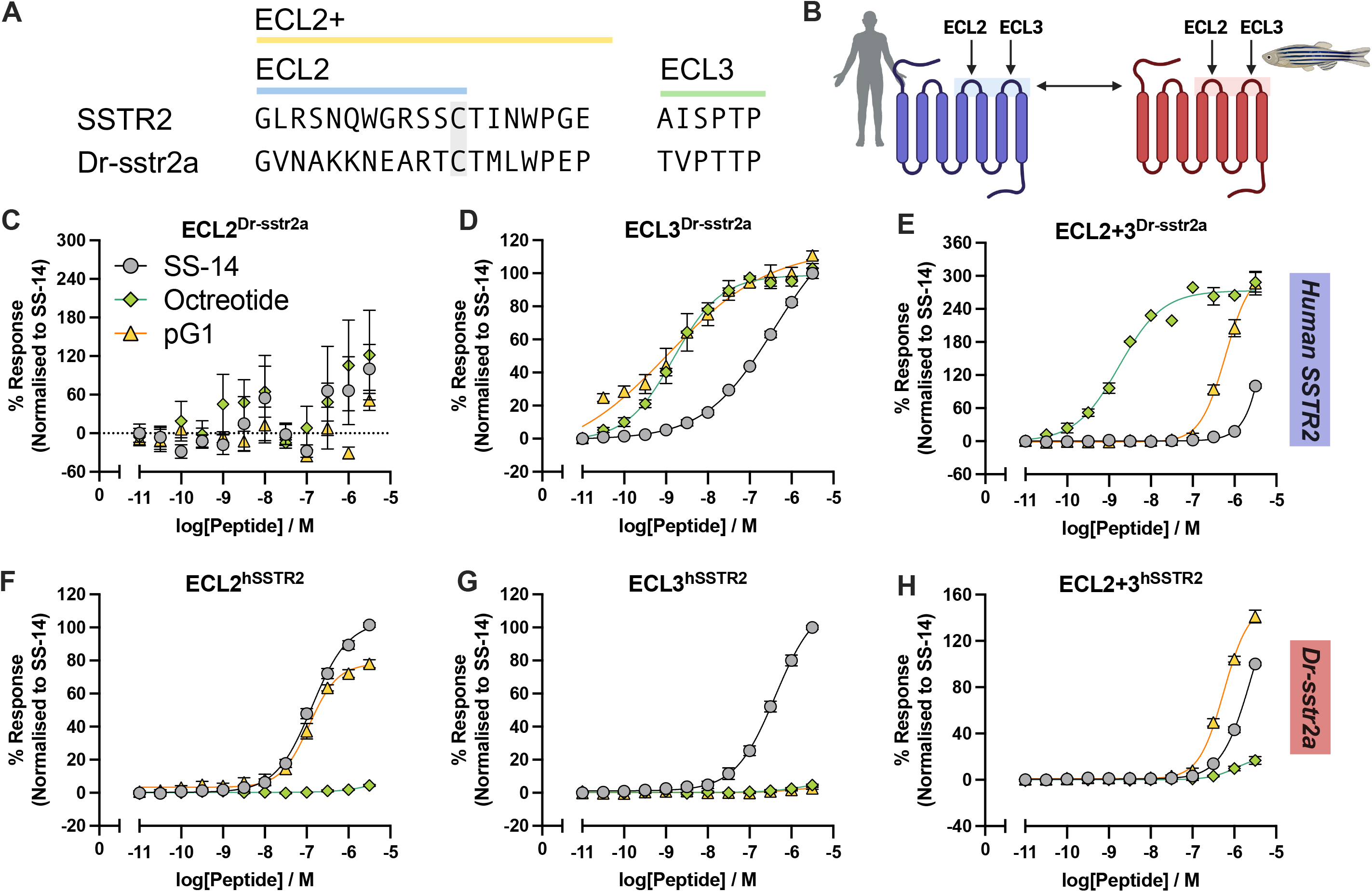
Extracellular loop swaps reveal differential dependence of pG1 and octreotide on ECL architecture. (A) Sequence alignment of the ECL2 and ECL3 regions of the human SSTR2 and Dr-sstr2a. (B) Diagram illustrating the design of human SSTR2 and Dr-sstr2a chimeras. (C-E) Representative concentration–response curves showing activation of human SSTR2 chimeras containing Dr-sstr2a ECL2, ECL3, or both ECL2 and ECL3 by SS-14, pG1 and octreotide. (F-H) Representative concentration–response curves showing activation of Dr-sstr2a chimeras containing human ECL2, ECL3, or both ECL2 and ECL3 by SS-14, pG1 and octreotide. All data are presented as mean ± S.E.M. from three independent biological replicates, each performed in technical duplicate.

### pG1 and octreotide differentially engage extracellular loops of SSTR2

We generated chimeric receptors in which ECL regions (ECL2, ECL3, or both) were exchanged between human SSTR2 and zebrafish Dr-sstr2a and assessed signaling using the PRESTO-Tango β-arrestin recruitment assay (21) **(Fig. 2A-B)**. For ECL2 swaps, substitutions were confined to the region between the conserved cysteine residues (denoted as ECL2 in **Fig. 2A**). Most chimeric receptors exhibited comparable cell surface expression with their wildtype receptors, except for hSSTR2 with Dr-sstr2a ECL3 and Dr-sstr2a with both ECL2 and ECL3 which showed a somewhat reduced cell surface expression (**Fig. S2**).

Substitution of zebrafish Dr-sstr2a ECL2 into human SSTR2 profoundly disrupted receptor signaling, as SS-14, octreotide and pG1 all failed to activate this chimera, indicating that the integrity of human ECL2 is essential for human SSTR2 function **(Fig. 2C)**. In contrast, replacement of human ECL3 alone did not impair activation by octreotide or pG1, while SS-14 potency was reduced by close to ten-fold relative to wild-type SSTR2 **(Fig. 2D)**. Combined replacement of both human ECL2 and ECL3 with zebrafish sequences differentially affected ligand responses: octreotide retained substantially higher potency than pG1 despite 50% less cell surface expression compared to wildtype SSTR2 **(Fig. S2)**, whereas SS-14 signaling was completely abolished **(Fig. 2E)**. These patterns indicate that the ECL regions contribute differently to signaling by SS-14, pG1, and octreotide. However, the observed effects may reflect not only ligand-specific receptor contacts, but also differences in ECL conformational dynamics and their influence on receptor activation.

Reciprocal substitutions in the zebrafish receptor supported differential contributions of ECL regions to pG1 and octreotide signaling. Chimeric Dr-sstr2a receptors containing human SSTR2 ECL residues showed reduced responsiveness to SS-14 **(Fig. 2F-H)**. Notably, introduction of human ECL2 into Dr-sstr2a partially restored pG1 responsiveness but did not confer responsiveness to octreotide (**Fig. 2F**). Introduction of human ECL3 alone was insufficient to enable activation by either ligand **(Fig. 2G)**, whereas introduction of both human ECL2 and ECL3 again partially enhanced pG1 activity, albeit with reduced potency relative to human SSTR2 **(Fig. 2H)**. Together, these results support a role for human ECL2 in pG1 responsiveness, while also suggesting that additional receptor features contribute to the full pharmacological response. To further delineate the structural basis of these interactions, we initially performed computational docking using AlphaFold2. However, the predicted models failed to reveal ligand-specific differences between octreotide- and pG1-bound SSTR2 (data not shown). We therefore pursued cryo-EM to resolve the pG1-SSTR2 complex.

### Structural comparison of pG1- and octreotide-bound SSTR2 reveals conserved core binding and broader extracellular engagement by pG1

We resolved the cryo-EM structure of pG1 bound to human SSTR2 in complex with Gαi3 at 1.9 Å resolution (pdb: 12LM) **(Fig. 3A, Fig. S3-4, Table S2)**. To compare ligand recognition modes, we superimposed the pG1–SSTR2 structure with the previously reported octreotide-bound SSTR2 complex (pdb: 7T11) (22) **(Fig. S5)**. The receptor conformation is highly similar between the two structures, with conserved rearrangements of the transmembrane core characteristic of active-state Class A GPCRs (22). In both complexes, the conserved ligand Lys side chain engages D122^3.32^ (22) (numbering according to the Ballesteros-Weinstein Scheme) (23), and the conserved D-Trp residue is packed into a shared hydrophobic and aromatic pocket formed by F208^5.38^ and F272^6.51^ (22, 24, 25) **(Fig. S5)**, underscoring a shared orthosteric anchoring mechanism.

**Figure 3:**
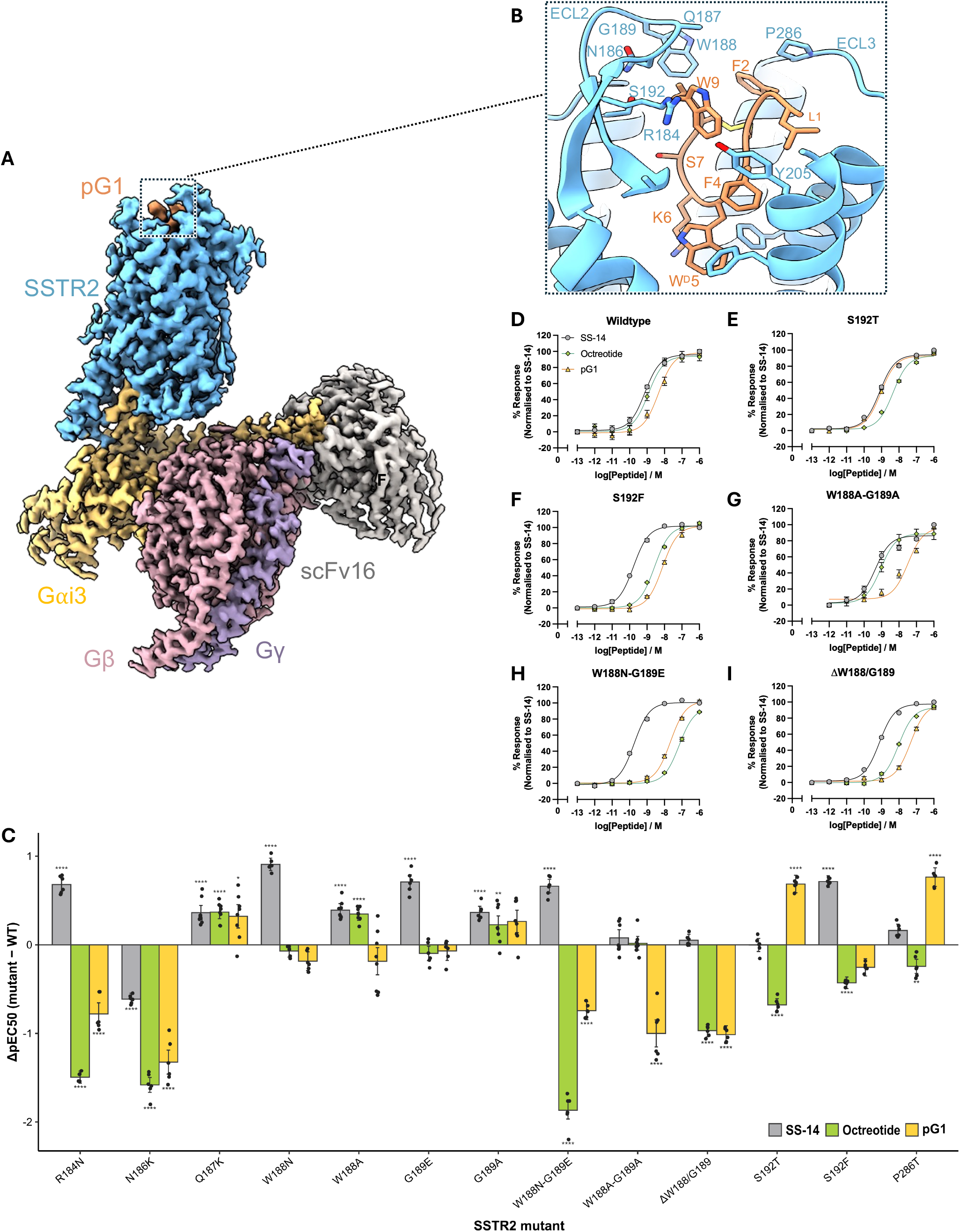
Subtle differential engagement of pG1 and octreotide with ECL2 revealed by cryo-EM structure determination and receptor mutagenesis. (A) Overall structure of human SSTR2 (blue) bound to pG1 (orange) in complex with Gαi3 (gray), Gβ (gold), Gγ (salmon), and scFv16 (mint). (B) Close-up views of the extracellular loop regions (ECL2 and ECL3) highlighting residues involved in pG1 engagement. (C) Bar plot summarizing changes in pEC_50_ values for activation of wild-type and mutant SSTR2 by SS-14, octreotide, and pG1 (plotted in mean ± S.D.). Changes were calculated by subtracting pEC50,wildtype from pEC50,mutants. Statistical significance was determined using one-way repeated-measures ANOVA followed by Dunnett’s post hoc test, comparing each mutant to the corresponding wild-type receptor for the same ligand (**p < 0.01 and ****p < 0.0001). (D-I) Representative concentration–response curves showing activation of SSTR2 wildtype and mutants by SS-14, pG1 and octreotide. All data are presented as mean ± S.E.M. from three independent biological replicates, each performed in technical duplicate.

Despite this conserved core engagement, the complexes show subtle differences in the positioning of the peptide termini relative to ECL2. Octreotide makes fewer contacts with ECL2 than pG1: its C-terminal Thr-ol8 forms a hydrogen bond with S192 and van der Waals contacts with R184, N186, W188, and its Phe3 packs against Y205^5.35^ near ECL2. In pG1, Trp9 makes additional, closer contacts with R184 and Q187 in addition to the ECL2 residues that interact with octreotide. Furthermore, Leu1 and Phe4 of pG1 pack against Y205^5.35^, while Phe2 of pG1 and D-Phe1 of octreotide both contact P286 on ECL3 **(Fig. 3B)**. These subtle differences in extracellular engagement are consistent with the distinct effects of ECL substitutions on pG1 and octreotide signaling. However, they do not exclude contributions from ligand-dependent ECL conformational dynamics to receptor activation.

### Receptor mutagenesis identifies ECL2 determinants of ligand recognition

To assess the functional contribution of ECL contacts identified in the cryo-EM structure, and to investigate the reduced pG1 responsiveness of Dr-sstr2a, we introduced single or double substitutions of human SSTR2 ECL2 and ECL3 residues with their Dr-sstr2a counterparts. We then measured SS-14, octreotide, and pG1 activity using the TRUPATH G protein dissociation assay (26). We selected this assay for further characterization as the receptor constructs used in the PRESTO-Tango assay are hybrids with C-terminal tail of the vasopressin receptor 2 to facilitate β-arrestin recruitments (21) whereas SSTR2 receptor constructs without C-terminal changes are used in the TRUPATH G protein dissociation assay (26). Surface expression was comparable across mutants, except for W188N/G189E which showed a strongly reduced expression **(Fig. S6)**. Effects are reported as ΔpEC_50_ relative to wildtype SSTR2.

SS-14 response to all receptor mutants was largely unaffected, exhibiting no more than one-log changes in potency **(Fig. 3C, Fig. S7)**. R184N and N186K reduced the potency of both octreotide and pG1, consistent with prior reports showing that alanine substitutions at N186 and R184 impair octreotide signaling **(Fig. 3C)** (22). S192T had opposite effects on the two peptides, enhancing pG1 potency while reducing octreotide activity **(Fig. 3D-E)**. Substitution with the bulkier residue S192F modestly reduced octreotide potency without a significant effect on pG1 **(Fig. 3F, Fig. S7)**. Q187K increased the potency of octreotide and pG1 to a similar extent **(Fig. S7)**. Finally, P286T in ECL3 enhanced pG1 potency by 0.8 log with only a small reduction in octreotide potency **(Fig. 3C, Fig. S7)**, consistent with the chimera experiments in which insertion of human ECL2 and ECL3 into Dr-sstr2a selectively increased pG1 responsiveness.

ECL2 undergoes ligand-induced rearrangement and contributes to peptide recognition and agonist potency at SSTR2 (22, 24, 25, 27). W188 and G189 sit at the tip of ECL2, where W188 packs against the C-terminal Thr-ol8 of octreotide (22). However, superimposing SSTR2 bound to octreotide and pG1 only reveals limited insight, so we used receptor mutagenesis to probe its contribution to ligand recognition. Single substitutions of W188 (W188N, W188A) or G189 (G189E, G189A) had little or no effect on pG1 or octreotide potency **(Fig. 3C, Fig. S7)**. However, ligand-specific effects emerged with combined substitutions. The alanine double substitution W188A/G189A decreased pG1 potency by 1.0 log with no effect on octreotide or SS-14 **(Fig. 3G)**. In contrast, W188N/G189E reduced octreotide potency to a greater extent by 1.9 log and pG1 potency by 0.7 log, although this mutant showed reduced surface expression **(Fig. 3H)**. Deletion of the W188/G189 tip (ΔW188/G189) reduced the potency of pG1 and octreotide equally (by approximately 1.0 log) **(Fig. 3I)**, indicating that the length and structural integrity of the ECL2 tip are required for both peptides.

Collectively, these results identify R184, N186 and the W188/G189 tip as ECL2 elements required for pG1 and octreotide activity, while S192, the W188/G189 side chains and P286 differentiate the activity of these two peptides. Octreotide is more sensitive to changes at S192 and to the polar W188N/G189E substitution, whereas pG1 depends on the W188/G189 side chains and is enhanced by S192T and P286T. These data support a model in which the two peptides engage the ECL network differently, and may explain why Dr-sstr2a hybrids with ECL2 from human SSTR2 partially restored pG1 potency **(Fig. 2F)**.

### Rational design of potent SSTR2-selective pG1–octreotide hybrid agonists

Guided by our structure showing ECL2 engagement by pG1, we aimed to improve pG1 potency by modifying residues predicted to contribute to ECL contacts while retaining SSTR2 selectivity. Because Trp9 of pG1 makes extensive contact at the ECL2, we first examined the contribution of aromatic character and stereochemistry at this C-terminal position. Potency was measured using the TRUPATH G protein dissociation assay (26) while receptor subtype selectivity was measured using PRESTO-Tango β-arrestin recruitment across SSTR1–5 (21).

Substitution of Trp9 with Phe (pG1[W9F]) did not significantly alter potency (pEC_50_ ± S.E.M: 8.91 ± 0.20, pG1[W9F] and 8.75 ± 0.10, pG1) **(Fig. 4A, Fig. S8, Table S3)**, whereas replacement with Tyr (pG1[W9Y]) reduced potency by 4-fold (pEC_50_ ± S.E.M: 8.12 ± 0.27). In contrast, inversion of stereochemistry at position 9 (L-Trp9 to D-Trp9, pG1[W^D^9]) reduced SSTR2 potency about 36-fold, demonstrating that the L-configuration of Trp9 is critical for ECL2 interaction **(Fig. 4A, Fig. S8, Table S3)**. Together, these data indicate that the aromatic side chain at position 9 is sensitive to substitution and stereochemical positioning at Trp9 are essential for optimal receptor engagement. None of these substitutions altered SSTR2 selectivity **(Fig. 4A, Fig. S9)**.

**Figure 4:**
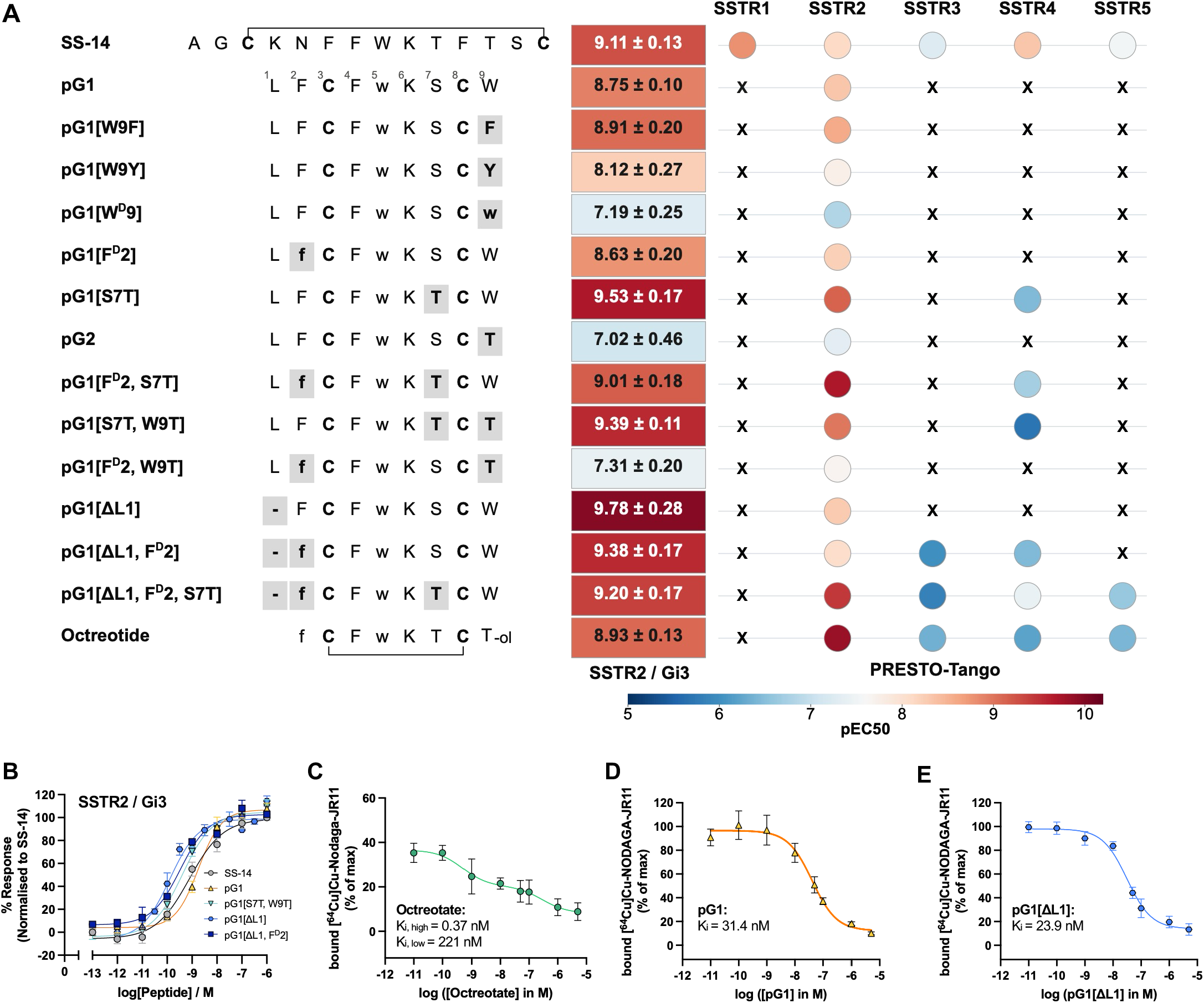
pG1–octreotide hybrid analogs exhibit enhanced potency. (A) Heatmap summarizing the potencies, measured at SSTR2 dissociating Gi3 protein, and receptor subtype selectivity of SS-14, pG1, octreotide measured using the TRUPATH G protein dissociation and PRESTO-Tango β-arrestin recruitment assays. Data are presented as mean ± S.E.M. from at least three independent biological replicates, each performed in technical duplicate. w, D-trp; –ol denotes a C-terminal alcohol modification. (B) Representative concentration–response curves showing SSTR2/Gi3 activation by SS-14, pG1 and pG1 analogs with improved potencies. All data are presented as mean ± S.E.M. from three independent biological replicates, each performed in technical duplicate. (C-E) Representative concentration–response curves showing binding affinities of octreotate, pG1 and pG1[ΔL1] respectively measured using a radiometric competitive binding assay with [64Cu]Cu-NODAGA-JR11 as the radiolabeled SSTR2 antagonist in mouse pheochromocytoma cells. All binding data are presented as mean ± S.D. from six independent biological replicates.

We next introduced octreotide-inspired substitutions into the pG1 backbone to test whether they could further enhance pG1 potency without losing SSTR2 selectivity. L-Phe2 to D-Phe2 (pG1[F^D^2]) changed neither potency nor selectivity (pEC_50_ ± S.E.M: 8.63 ± 0.20). In contrast, substitution of Ser7 with Thr (pG1[S7T]) increased potency by 6-fold (pEC_50_ ± S.E.M: 9.53 ± 0.17), but also introduced SSTR4 activation **(Fig. 4A, Fig. S8-9)**. Consomatin pG2, a variant of pG1 derived from another *C. geographus* venom peptide that contains Thr9 (28), was 54-fold less potent than pG1 (pEC_50_ ± S.E.M: 7.02 ± 0.46) but remained SSTR2-selective. Because pG2 retains Ser7 and stays selective, whereas S7T gains SSTR4 activity, Ser7 appears to contribute to SSTR2 subtype discrimination.

We next evaluated combinatorial substitutions. The double mutant pG1[F^D^2, S7T] was 2-fold more potent (pEC_50_ ± S.E.M: 9.01 ± 0.18) than pG1, but similar to pG1[S7T], also activated SSTR4 **(Fig. 4A, Fig. S8-9)**. The double mutant pG1[S7T, W9T] exhibited 4-fold increase in potency (pEC_50_ ± S.E.M: 9.39 ± 0.11) and showed weaker SSTR4 activation than pG1[S7T] **(Fig. 4A-B)**. In contrast, combining D-Phe2 with Thr9 (pG1[F^D^2, W9T]) reduced potency by 28-fold while remaining SSTR2-selective **(Fig. 4A, Fig. S8-9)**, further suggesting retaining Ser7 is important for SSTR2 selectivity.

Finally, we evaluated the effect of truncating the N-terminus, as observed in octreotide. Truncation of the N-terminal Leu1 (pG1[ΔL1]) produced the most potent analog identified in this study (pEC_50_ ± S.E.M: 9.78 ± 0.28), about 11-fold more potent than pG1, while remaining SSTR2-selective **(Fig. 4A-B, Fig. S8-9)**. This suggests that removal of Leu1 may alleviate steric or conformational constraints and improve engagement of pG1 with the extracellular R184-Y205^5.35^ network **(Fig. 3B)**. Introducing D-Phe2 (pG1[ΔL1, F^D^2]) alone or in combination with the Thr7 substitution (pG1[ΔL1, F^D^2, S7T]) resulted in 4.3- and 2.8-fold increases in potency, respectively **(Fig. 4A-B, Fig. S8-9)**, however, neither exceeded the potency of pG1[ΔL1]. These truncated combinations also lost selectivity: pG1[ΔL1, F^D^2] additionally activated SSTR3 and SSTR4, and pG1[ΔL1, F^D^2, S7T] activated SSTR3, SSTR4 and SSTR5, resembling the profile of octreotide **(Fig. 4A, Fig. S9)**. Thus, pG1[ΔL1] offers the best balance of potency and SSTR2 selectivity.

### pG1 and pG1[ΔL1] retain binding to endogenous mouse SSTR2

To complement human SSTR2 receptor potency and selectivity studies, we characterized the binding affinities of the two most potent and SSTR2-selective peptides identified in this study, pG1 and pG1[ΔL1]. We performed a radiometric competitive binding assay using the radiolabeled SSTR2 antagonist [^64^Cu]Cu-NODAGA-JR11 (29, 30) in mouse pheochromocytoma cell (MPC), which endogenously express high levels of SSTR2 (31, 32). Octreotate, an octreotide analog bearing Thr8 instead of Thr-ol8, was included as reference ligand. Although the use of endogenous mouse SSTR2 precludes a direct quantitative comparison with activation assays performed using overexpressed human SSTR2, it provides complementary evidence of ligand binding in a native cellular receptor context and supports the utility of these ligands in mouse models of diseases associated with SSTR2.

Octreotate displayed a biphasic competitive binding curve, with *K*_i_ values of 0.37 nM and 221 nM for the high- and low-affinity binding sites, respectively **(Fig. 4C)**. In contrast, pG1 displaced the radioligand in a monophasic manner (K_i_: 31.4nM; p*K*_i_ ± S.E.M.: 7.50 ± 0.05) **(Fig. 4D)**. Removal of Leu1 produced only modest improvement in binding affinity (K_i_: 23.9 nM; p*K*_i_ ± S.E.M.: 7.62 ± 0.02) **(Fig. 4E)**. pG1[W^D^9], the least potent pG1 analog tested in the G protein dissociation assay, also showed reduced binding affinity (K_i_: 79.8 nM; p*K*_i_ ± S.E.M.: 7.10 ± 0.05) **(Fig. S10)**. This suggests that the binding affinities of these analogs are consistent with their functional potencies. Thus, in this endogenous mouse SSTR2 system, the increased activity of pG1[ΔL1] is not accompanied by a commensurate affinity gain, suggesting that the potency difference may reflect either more efficient receptor activation or G protein coupling, or species-specific differences between mouse and human SSTR2.

## Discussion

Venom peptides have been shaped by millions of years of evolution into ligands with high potency, selectivity, and metabolic stability (33, 34). These properties enable efficient prey capture and predator deterrence and have established venom peptides as a valuable source of pharmacological probes and therapeutic leads (35–37). Early efforts focused on cone snail toxins that target ion channels (38), but it has become increasingly clear that certain venom components also modulate GPCRs (39), expanding their relevance to GPCR peptide ligand discovery (33, 40). Our recent identification of consomatins, cone snail venom peptides that selectively target human SSTR subtypes (28), has opened new opportunities for developing venom-inspired subtype-selective ligands for SSTR–associated diseases. We previously showed that consomatin Fj1 from *Conus fijisulcatus* potently and selectively activates the SSTR4 and provides analgesia in several animal models of pain (28). Structure-activity-guided optimization of Fj1 yielded the most potent and selective SSTR4 peptide agonists reported to date (41). Here, we extend this work to consomatin pG1 from *Conus geographus*, leveraging its high potency and selectivity for human SSTR2, lack of activity at the fish receptor, and cryo-EM structural analysis to define the ligand and receptor determinants of SSTR2 activation and guide rational analog design.

Despite strong conservation of orthosteric-site residues between human SSTR2 and zebrafish Dr-sstr2a, pG1 failed to activate the zebrafish receptor, providing a useful tool for identifying determinants of activity outside the orthosteric binding pocket. Consistent with this interpretation, exchanging ECL regions demonstrated their functional importance. Incorporation of human ECL2 into Dr-sstr2a conferred pG1 responsiveness, whereas replacement of human SSTR2 ECLs with zebrafish sequences abolished it. In contrast, octreotide remained active at human SSTR2 chimeras bearing zebrafish ECL2 and ECL3, indicating that its activity is less sensitive to ECL composition. Thus, ECL divergence, rather than differences in the conserved orthosteric pocket, likely underlies the species-selective loss of pG1 activity and illustrates that orthosteric conservation alone may be insufficient to predict pharmacological equivalence between receptor homologs.

To further resolve these ECL interactions, we defined the molecular structural interactions by solving the pG1-bound human SSTR2 cryo-EM structure. pG1 preserves the conserved D-Trp-Lys pharmacophore characteristic of octreotide. Both pG1 and octreotide anchor within the canonical orthosteric pocket through conserved interactions, including a salt bridge with D122^3.32^ and π-stacking of D-Trp with F208^5.38^ and F272^6.51^ (22). They diverge at the C-terminus: Trp9 of pG1 forms a broader contact network with ECL2, including R184 and Q187, than Thr-ol8 of octreotide. The C-terminus of the minimized cyclic scaffold thus engages ECL2 extensively, identifying this region as a key for tuning the ligand–ECL2 interface.

Receptor mutagenesis provided further support for a role of ECL2 in ligand-specific activation. Consistent with prior reports (22), substitution of residues at the base of ECL2, including R184 and N186, with their Dr-sstr2a counterparts broadly impaired signaling by all ligands. In contrast, mutations at the ECL2 tip, W188 and G189, produced more ligand-dependent effects. Their simultaneous substitution with alanine disproportionately reduced pG1 activity relative to octreotide. However, replacement with the zebrafish residues or deletion of this segment markedly reduced the potency of both peptides, highlighting the broader importance of the ECL2 tip for ligand recognition and activation. Together, these data indicate that pG1 and octreotide depend on a largely shared set of ECL2 residues, while differences at the ECL2 tip contribute to pG1’s distinct extracellular engagement and to the loss of activity at Dr-sstr2a.

Guided by these structural and receptor mutagenesis insights, we next engineered pG1– octreotide hybrid analogs to explore if such hybrids could provide enhanced SSTR2 potency. Substitutions at the C-terminal Trp9 residue with other aromatic residues demonstrated that changing the C-terminal end does not affect receptor selectivity but does impact potency based on side-chain hydrophobicity. By contrast, replacing Ser7 within the cysteine loop with the octreotide-equivalent Thr shifted subtype selectivity, likely through altered contacts with S192 and T194 in ECL2. Truncating the N-terminus to match the octreotide loop size gave pG1[ΔL1], the most potent analog in the series, consistent with relief of a steric clash between Leu1 and Y205^5.35^. Selected C-terminal modifications can improve potency while preserving selectivity, whereas the core pharmacophore and N-terminus govern subtype preference. Together with their high potency and selectivity at human SSTR2, the measurable binding of pG1 and pG1[ΔL1] to endogenously expressed mouse SSTR2 supports their utility as human SSTR2-targeted probes and potential therapeutic leads, while also enabling their future evaluation in mouse models of disease.

Our studies have several limitations. First, although the introduction of human ECL2 into Dr-SSTR2a partially restored pG1 potency, it did not fully recapitulate the potency observed at wild-type human SSTR2. This may reflect the contribution of additional receptor determinants, potentially including residues within the transmembrane core, as well as the structural integrity and conformational context of ECL2 within the full receptor scaffold. These factors may contribute to pG1-mediated SSTR2 activation and remain to be investigated.

In addition, the distinct competitive binding profiles of octreotate, pG1, and pG1[ΔL1] are not yet fully explained. Octreotate exhibited a biphasic competition curve, consistent with binding to receptor populations in free and G protein-bound active states (42), whereas pG1 and pG1[ΔL1] displayed monophasic curves. These patterns may reflect different preferences for receptor conformations or G protein-coupled states, but the underlying mechanisms remain unclear. The endogenous G protein complement of MPC cells has not been characterized, and the TRUPATH G protein dissociation assay was limited to Gi3 in line with our resolved cryo-EM structure. Future studies should therefore define the endogenous G protein composition of MPC cells and determine how G protein coupling influences the binding profiles and pharmacological properties of these peptides. Finally, the distinct C-terminal chemistries of pG1 and octreotate, including a free carboxylate and a threoninol group, respectively, may also contribute to their differential receptor interactions and binding profiles (43). This possibility remains to be tested experimentally.

In summary, our findings reinforce and extend emerging evidence that ECL architecture is a key determinant of SSTR2 ligand selectivity and efficacy (22, 25, 27, 44–46). While the conserved transmembrane core provides a common anchoring framework for somatostatin analogs, divergence within ECL2 modulates ligand-receptor interaction, species selectivity, and activation efficacy. Exploiting these extracellular interactions offers a generalizable strategy for developing next-generation, subtype-selective SSTR2 therapeutics and emphasizes the continued value of venom-derived peptides as templates for rational GPCR drug design.

### Experimental procedures

#### Chemical reagents

Unless otherwise specified, all chemical and cell culture reagents were sourced from Sigma and ThermoFisher respectively. SS-14 was obtained from GenScript. pG1, pG2 and octreotide were custom synthesized by Genscript. All pG1 analogs were synthesized in house using solid phase peptide synthesis (SPPS) as detailed in “Peptide synthesis”. The mass and purity (> 95 %) of the peptides were verified by RP-HPLC and MS prior to use (Supporting Information, under ‘General Peptide Synthesis’ section).

#### Cell culture

HEK293T cells were given by Dr. Justin English (University of Utah, US) and were cultured in DMEM supplemented with 10% Fetal Bovine Serum (Gibco) and 100 U/mL penicillin and 100 µg/ml streptomycin at 37°C in a humidified incubator supplemented with 5% CO_2_. HTLA cells were given by Prof. Bryan Roth (University of North Carolina, Chapel Hill, US) and were maintained in Dulbecco’s modified Eagle’s medium (DMEM) supplemented with 10% Fetal Bovine Serum (FBS) (Biowest) and 100 U/mL penicillin and 100 µg/mL streptomycin at 37°C in a humidified incubator supplemented with 5% CO_2_. Constant selective pressure for tTA-dependent luciferase reporter and a β-arrestin 2 tobacco etch virus protease fusion gene were maintained with the addition of 100 µg/mL hygromycin B and 2 µg/ml puromycin (Tango growth medium). MPC cells (clone 4/30PRR (47)) were routinely cultured in collagen-coated flasks as described previously (31). Mycoplasma testing was conducted every 3 months to ensure the lack of mycoplasma contamination.

#### Plasmids

Tango constructs for human SSTR1-5 were kind gifts from Prof. Bryan Roth (University of North Carolina, Chapel Hill, USA) (Addgene kit # 1000000068). Tango constructs for zebrafish Dr-sstr2a (sequence accession number: XP_005170178.1) and Dr-sstr2b (sequence accession number: XP_017210358.1), and Tango constructs used in the ECL swap experiments were custom-synthesized by Twist Biosciences using the Tango construct template as described (21). 3xHA-tagged human SSTR2 construct was obtained from cDNA.org (cDNA resource center, Bloomsburg PA, US). TRUPATH G protein constructs were gifts from Prof. Bryan Roth (University of North Carolina, Chapel Hill) (Addgene kit #1000000163) (26). SSTR2 mutants (i.e. R184N, N186K, Q187K, W188N, W188A, G189E, G189A, W188N-G189E, W188A-G189A, ΔW188/G189, S192T, S192F, P286T) inserted in pcDNA3.1-N-terminal tagged HA used in this study was custom synthesized by GenScript. All DNA constructs were sequenced and confirmed prior to use.

#### PRESTO-Tango β-arrestin recruitment

The assay was done as described in (19). Briefly, HTLA cells (1.2 million cells/well) were seeded in 6-well plates in Tango growth medium and incubated overnight. On day 2, medium was changed to DMEM supplemented with 10% FBS without hygromycin B and puromycin. Cells were then transfected using Polyfect (Qiagen) according to manufacturer’s protocol. On day 3, 15,000 cells in 40 µL DMEM supplemented with 1 % dialyzed FBS per well were seeded in poly-D-lysine-coated white clear-bottom 384 well plates (Corning) and incubated overnight. On day 4, the Tango assay medium was changed and 10 µL of each of the peptide was added at 5x final concentration. The peptides were resuspended in HBSS with 20 mM HEPES, 1 mM of CaCl_2_, 1 mM of MgCl_2_, pH adjusted to 7.4 with 1 M NaOH, which was supplemented with 0.1 % bovine serum albumin (BSA) (Tango assay buffer). After overnight incubation, on day 5, the medium and the compounds were removed from the cells and 20 µL of 1:20 dilution of BrightGlo (Promega) in Tango assay buffer supplemented with 0.01 % pluronic F68 was added to the cells. Plates were incubated for 20 minutes in the dark at room temperature. Luminescence was measured on PherastarFX (BMG Labtech) with each well integrated for 1 second.

#### Expression and Purification of Somatostatin receptor 2

Pellets containing SSTR2 were expressed in Sf9 cells and snap-frozen in liquid nitrogen for storage. To begin purification of the receptor, cell pellets were thawed and lysed in hypotonic lysis buffer containing 10 mM HEPES pH 7.5, Pierce Universal Nuclease, protease inhibitor cocktail, 1 mM benzamidine, 1 mM MgCl_2_, and 1 mM EDTA. The solution was suspended on a stir plate for 1 hour at 4°C. The membranes were harvested through ultracentrifugation at 100,000xg for 35 minutes. Pellets were resuspended in a solubilization buffer containing 20 mM HEPES at pH 7.5, Pierce Universal Nuclease, 500 mM NaCl, protease inhibitor cocktail, 1 mM benzamidine, 10% glycerol, and 1 mM MgCl_2_. Subsequently, detergent was added dropwise to the resuspended pellets to reach a final concentration of 1% LMNG/0.2% sodium cholate/0.2% CHS while stirring at 4°C. Following gentle stirring at 4°C for 3 hours, the solution was subjected to ultracentrifugation at 100,000xg for 35 minutes to remove insoluble debris. Supernatant was then pooled and supplemented with 20 mM imidazole before being loaded over a cobalt resin column that had been washed with 10 column volumes of buffer containing 20 mM HEPES at pH 7.5, 500 mM NaCl, 20 mM imidazole, and 0.1% LMNG/0.01% CHS. Protein was eluted from the column with 2 column volumes of buffer containing 20 mM HEPES pH 7.5, 250 mM NaCl, 10% glycerol, 250 mM imidazole, and 0.01% LMNG/0.001% CHS. Purified SSTR2 was concentrated via microcentrifugation then loaded over a Superdex 200 column in buffer containing 100 mM NaCl, 20 mM HEPES pH 7.5, 0.01% LMNG/0.001% CHS, and 5% glycerol to be subjected to SEC chromatography. Peak fractions were collected and pooled together, then concentrated. Supplementation with 5% glycerol was followed by snap-freezing with liquid nitrogen for storage and use with complexation.

#### Expression and Purification of Dominant Negative Gi3

To start purifying dominant negative Gi3, cell pellets expressed in Hi5 cells (infected with viruses for Gαi3, Gβγ, and ric8a) were thawed and resuspended in lysis buffer containing 20 mM HEPES pH 7.5, 5% glycerol, 1 mM MgCl_2_, 1 mM EDTA pH 8, 100 µM GDP, 5 mM Beta-Mercaptoethanol, protease inhibitor cocktail, and Pierce Universal Nuclease, then gently at 4°C. Lysate was then subjected to ultracentrifugation at 100,000xg for 35 minutes. Supernatant was collected and supplemented with 30 mM imidazole, then loaded over Ni-NTA columns and allowed to flow through. Subsequently, the columns were washed with 10 column volumes of a series of buffers containing 30 mM imidazole and 50% solubilization/50% E2 buffer, 25% solubilization/75% E2 buffer, 12.5% solubilization/87.5 E2 buffer, and 100% E2 buffer (E2 buffer contained 20 mM HEPES pH 7.5, 5% glycerol, 100 mM NaCl, 1 mM MgCl_2_, 0.05% LMNG/0.005% CHS, 5 mM Beta-Mercaptoethanol, and 100 µM GDP). Purified protein was eluted with 3 column volumes of E2 buffer and 250 mM imidazole, then incubated with 1 mg of 3C protease per 50 mg of protein with overnight dialysis in a 3,500 MWCO dialysis bag at 4°C. The following day, the Ni-NTA column was washed with 10 column volumes of E2 buffer and 30 mM imidazole. Overnight cleavage was loaded over the column and flow through was collected. The column was washed with 2 additional column volumes of E2 buffer and 30 mM imidazole, and flow through was collected. The purified protein was concentrated to 500 µL in a 30 kDa MWCO spin concentrator and loaded over a Superdex 200 column in buffer containing 20 mM HEPES pH 7.5, 5% glycerol, 1 mM MgCl_2_, 100 mM NaCl, 100 µM TCEP, 20 µM GDP, and 0.01% LMNG/0.001% CHS for SEC chromatography.

#### Expression and Purification of scFv16

scFv16 was purified as described previously (48, 49). Purified scFv16 was then concentrated to 13 mg/mL and snap-frozen with liquid nitrogen in a buffer containing 20 mM HEPES pH 7.5, 100 mM NaCl, and 15% glycerol for storage and future use.

#### Formation and Purification of SSTR2-Gi3-scFv16 Complex

SSTR2 and peptide were incubated on ice for 1 hour. Complex formation began by mixing receptor with molar excess of wildtype Gi3 in a buffer containing 20 mM HEPES pH 7.5, 100 μM TCEP, 100 μM LFT, 100 mM NaCl, and 0.01% LMNG, then incubated for 1 hour on ice. scFv16, apyrase, and HRV-3C protease were added, followed by overnight incubation on ice. The complex was diluted with buffer of 5-fold more volume consisting of 20 mM HEPES pH 7.5, 100 μM scFv16, 100 mM NaCl, and 0.01% LMNG/0.001% CHS. Diluted complex was loaded over an M1 flag resin column, then washed with buffer containing 20 mM HEPES pH 7.5, 10 μM scFv16, 100 mM NaCl, and 0.004% LMNG/0.0004% CHS. SSTR2-Gi3-scFv16 complex was eluted from the column with a buffer containing 20 mM HEPES pH 7.5, flag peptide, 50 μM scFv16, 10% glycerol, 100 mM NaCl, and 0.002% LMNG/0.0002% CHS. Eluent was supplemented with 100 μM TCEP then concentrated to a lower volume before being loaded onto a Superdex 200 column with buffer of 20 mM HEPES pH 7.5, 100 mM NaCl, 100 μM scFv16, 100 uM TCEP, 2mM MgCl_2_, and 0.001% LMNG/0.0001% CHS/0.00033% GDN for SEC chromatography. Peak fractions were pooled and collected for cryo-EM analysis.

#### Cryo-EM Sample Prep and Imaging

Cryo-EM samples were prepared at 100% humidity and 4°C with a Vitrobot Mark IV. Sample was applied at 3 μL to UltrAuFoil R1.2/1.3 300 mesh grids that had been glow discharged in a Pelco unit for 45 seconds at 10 mA, followed by plunge freezing in liquid ethane. Grids were imaged on a G4 Titan Krios at 300 kV with a K3 direct electron detector and a Bioquantum energy filter. Details of the imaging parameters can be found in Table S2.

#### Data Processing and Model Building

All data was processed in cryoSPARC (50) with patch motion correction, patch CTF estimation, template-based picking with existing GPCR-G-protein complex templates, particle extraction, and a single round of 2D classification to remove junk particles. Further curation was performed with iterative rounds of *ab-initio* model generation with 6 classes followed by heterogeneous refinement. The final particle stack was used in an initial non-uniform refinement, followed by CTF refinement and reference-based motion correction. These particles were then treated with a final non-uniform and receptor/Ras domain local refinement. Initial models came from PDB:6OSA and 6OS9 for most components (51) while dominant negative Gi3 came from PDB:7T10 (22). Model building was performed in Coot (52) (with Ramachandran and torsion restraints applied) with iterative rounds of real space refinement in Phenix (53) (with secondary structure restraints applied). Details of the final maps and models can be found in Table S2.

#### TRUPATH G protein dissociation

The following protocol was adopted from (26). In brief, HEK293T cells were seeded onto 6-well plates at a density of 900,000 cells/well. Cells were transfected the next day with 3xHA-tagged human SSTR2 or SSTR2 mutants (0.3 µg), RLuc8-tagged G protein constructs (0.2 µg), pcDNA3.1-Gβ_3_ (0.2 µg), pcDNA-Gγ_2_-GFP2 (0.2 µg) (at a ratio of 1.5:1:1:1 and the total DNA amount per well was 0.9 µg) using using Polyfect (Qiagen) according to manufacturer’s protocol. 16 to 24 hours post-transfection, HEK293T cells were seeded onto white 96 well plate (PerkinElmer) coated with poly-L-lysine (PLL) at a density of 40,000 cells/well and were allowed to grow overnight. On the day of assay, cells were washed once with phosphate-based saline (PBS) and 80 µL of assay buffer which composed of PBS with 0.5 mM MgCl_2_, adjusted to pH 7.4, were added onto each well. 10 µL of Nluc luciferase substrate Prolume Purple (Nanolight coorperate) were added onto each well and incubated for 5 minutes. 10 µL of 10x concentrated ligands were then added manually onto the plate and BRET signal was measured instantaneously using PherastarFX (BMG technology) equipped with BRET1 410 (20 nm bandwidth) and 535 (25 nm bandwidth) filters continuously for 30 min.

#### BRET ratio calculation

BRET1 ratio was determined by dividing the BRET signal at 510 nm over that at 475 nm. The net BRET ratio was determined by subtracting the BRET ratio with drugs with the BRET ratio of vehicle control. Net BRET ratio over time was then plotted using GraphPad Prism (v.10.1.1). Net BRET ratio after 30-minutes ligand stimulation at each ligand concentration tested were determined and plotted against log(concentration) of drugs tested. Results of test ligands were normalized as percentage of maximal SS-14 response. Concentration response curves were fitted with a 4-parameter logistic equation (equation 1) using GraphPad Prism (v.10.1.1):

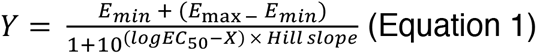

#### Receptor cell surface expression

HTLA or HEK293T cells were seeded onto 6-well plates at a density of 1.2 million cells/well or 900,000 cells/well respectively. The following day, cells were transfected with Tango-SSTR constructs (1.5 µg), 3xHA-tagged human wildtype SSTR2 or SSTR2 mutants (0.3 µg) or pcDNA3.1 (mock control), using Polyfect (Qiagen) according to manufacturer’s instructions. 16 to 24 hours post-transfection, transfected cells were seeded onto black 96 well plate (PerkinElmer) coated with PLL at a density of 40,000 cells/well and were allowed to grow overnight. For cell surface expression analysis, plates were washed with PBS, and cells were fixated with 4% paraformaldehyde (PFA) for 30 minutes on ice. Following fixation, cells were washed with PBS, and blocked with blocking buffer consisting of PBS supplemented with 0.1% BSA. Cells were incubated with either mouse monoclonal anti-HA antibody (ThermoFisher #26183) for detection of SSTR2 mutants, or mouse monoclonal M1-anti-FLAG-antibody (NovusBiologicals # NBP1-97410) for detection of Tango constructs for one hour on ice. After washing, cells were incubated with HRP-conjugated goat anti-mouse IgG, polyclonal secondary antibody (ThermoFisher #31430). Following washes, SuperSignal™ ELISA Femto Substrate (ThermoFisher, #37075) was added according to the manufacturer’s instructions, and chemiluminescence was measured.

#### Competitive binding assay

A number of 7.5×10^4^ MPC cells/well were seeded in collagen-coated 24-well microplates (CELLSTAR 24 Well Cell Culture Multiwell Plates, Polystyrene, Greiner bio-one, Item No. 662160) and grown for 4-5 days. For the competitive binding assays, cell culture medium was removed and replaced by fresh medium supplemented with the radioligand [^64^Cu]Cu-NODAGA-JR11 (2 nM, A_m_=25 MBq/nmol, prepared as previously described) (32) and increasing concentrations (0.01-5,000 nM) of competitor (final sample volume 0.5 mL). Samples were incubated for 60 min at 37°C. Incubation was stopped by washing with ice-cold PBS containing 0.9 mM CaCl_2_ and 0.5 mM MgCl_2_. Cells were lysed with 0.1 M NaOH containing 1% (w/v) SDS. Activity was measured in cell homogenates using the gamma counter Wizard (PerkinElmer). Protein content of cell homogenates was measured at A_280 nm_ (setting 1 Abs = 1 mg/mL) using a nanodrop spectrophotometer (Thermo-Fisher Scientific). For each competitor, two experiments (each in triplicate) were performed. The measured activity values for the cell homogenates were normalized to the activity values for cell homogenates (mean of six values) obtained in the absence of competitor. Plots of “bound [^64^Cu]Cu-NODAGA-JR11 (% of max)” as a function of competitor concentration (logarithmic scale) were each comparatively analyzed (Akaike’s Information Criterion) according to the models of “One site – Fit Ki” and “Two sites – Fit Ki” as implemented in GraphPad Prism (Version 11.0.2). “HotNM” (equals to radioligand concentration) was constrained to 2 (nM) and “HotKdNM”, “HotKdNMLo”, and HotKdNMHi” were constrained to 6.303 (nM, *K*_d_ of [^64^Cu]Cu-NODAGA-JR11 as determined by saturation binding analysis using intact MPC cells). For each competitor, the model with the higher probability was finally selected.

#### Statistical analysis

Data analyses were performed in GraphPad Prism (v.10.2.3). Statistical analysis was performed using one-way ANOVA with Dunnett’s test for SSTR2 mutant studies. Statistical significance was taken as p < 0.05.

#### Chemical synthesis

All solvents [*N,N*-dimethylformamide (DMF), dichloromethane (DCM), acetonitrile (ACN) and diethyl ether (Et_2_O) (Fisher Scientific)] and reagents [piperidine, *N*-methylmorpholine (NMM), *N*,*N*-diisopropylethylamine (DIPEA), trifluoroacetic acid (TFA), triisopropylsilane (TIPS), Fmoc-D-Phe-OH, Fmoc-D-Trp(Boc)-OH, (Sigma Aldrich), hexafluorophosphate azabenzotriazole tetramethyl uronium hydroxybenzotriazole (HATU) (Oakwood chemicals), sodium bicarbonate (NaHCO_3_), (Fisher Scientific) and Fmoc-D-Cys(SIT)-OH (Iris Biotech GmbH)] were purchased and used directly. Standard Fmoc amino acids (PurePep®) were purchased from Gyros Protein Technologies and used directly. 2-chlorotrityl chloride resin was purchased from Advanced Chemtech (cat. no. SC5054) and the first amino acid was manually coupled to the resin under standard conditions. Water (H_2_O) was obtained from a Milli-Q water purification system (Millipore) for preparative HPLC. 0.1% formic acid in water and 0.1% formic acid in acetonitrile (Fisher Scientific) were used for LC-MS.

#### HPLC and LC-MS

All crude peptides were purified with a 0.1% TFA buffered H_2_O (buffer A) and 0.1% TFA buffered 9:1 ACN:H_2_O (buffer B) gradient on an Agilent 1260 Infinity II HPLC system using a Phenomenex Jupiter C12, 4 µm (250 x 21.2 mm) (prod. numb. 00G-4396-P0-AX) preparative column at 15 mL.min^-1^. Fractions collected from HPLC were analyzed by LC-MS on a Phenomenex Jupiter C12, 4 μm (50 × 2 mm) (prod. numb. 00B-4396-B0) column at 0.5 mL.min^-1^ with a 0.1% formic acid buffered H_2_O (buffer A) and 0.1% formic acid buffered 9:1 ACN:H_2_O (buffer B) gradient on an Agilent 1260 Infinity II Quadrupole LC-MS system. Fractions containing targeted product (based on LC-MS) were collected and lyophilized. Purity of isolated peptides were determined by peak integration of blank subtracted HPLC spectra.

#### HRMS

High resolution mass spectrometry (HRMS) data was collected using an Agilent 1290 UHPLC. Mass analysis was performed using a Sciex 7600 Zeno-ToF.

#### General Peptide Synthesis

The synthesis of Consomatins and other related somatostatin-like peptides is described as follows. Standard Fmoc/tBu-based solid-phase peptide synthesis (SPPS) was used to synthesize all peptides. Peptides were synthesized on a Gyros PurePep® Chorus peptide synthesizer on a 25 µmol scale. The linear peptides were constructed on resins yielding either C-terminal acids (2-Chlorotrityl chloride resin) or C-terminal amides (TentaGel® R Ram resin). Deprotection of Fmoc was achieved with 20% piperidine in DMF for 2 min twice. Amide coupling was achieved by using 6 equivalents of amino acid (200 mM in DMF), 6 equivalents of HATU (200 mM in DMF) and 12 equivalents of NMM (600 mM in DMF) for 25 min. Side-chain protecting groups for amino acids were as follows: Trp and Lys, tert-butyloxycarbonyl (Boc); Ser and Thr, tert-butyl ether (tBu); Cys, trityl (Trt), sec-isoamyl mercaptan (SIT).

##### General procedure for disulfide bond formation

Cys(Trt) (trityl) was incorporated as the C-terminally directed cysteine, while Cys(SIT) (sec-isoamyl mercaptan) was incorporated as the N-terminally directed cysteine. Cleavage of the linear peptides was achieved by treatment of the peptidyl-resin with TFA/H_2_O/TIPS (TFA: trifluoracetic acid, TIPS: triisopropylsilane) (95:2.5:2.5) (4 mL). Formation of the intramolecular disulfide bond was accomplished by dissolution of the crude peptide in aqueous acetonitrile (ACN) (∼5 mL) and treatment of the resulting solution to pH 8 with 100 mM aqueous sodium bicarbonate (NaHCO_3_) solution. At this pH, the resulting C-terminally directed thiolate rapidly reacts with the N-terminally directed SIT-protected cysteine, generating the desired disulfide bond with concurrent liberation of isoamyl mercaptan as a byproduct. The reaction was monitored by LC-MS and was typically complete within 15 min. The resulting mixture was acidified to pH ∼2 and lyophilized prior to purification by preparative RP-HPLC.

## Data availability

Cryo-EM structure of pG1 bound to human SSTR2 in complex with Gαi3 is available through the Protein Data Bank (PBD) under accession number 12LM.

## Supporting information

This article contains supporting information.

## Acknowledgements

The technical assistance of Natalie Brenner and Andrea Suhr regarding peptide synthesis and the radiometric competitive binding assay is greatly acknowledged. MPC cells were kindly provided by Arthur Tischler. Furthermore, the authors wish to thank Dr. Reik Löser for discussions on the binding affinity data and for encouraging on establishing a new scientific collaboration.

## Funding and additional information

H.S.-H acknowledges funding from the National Institutes of Health (R01DK139317). MJR is a CPRIT Scholar in Cancer Research supported by CPRIT award RR230042. The content is solely the responsibility of the authors and does not necessarily represent the official views of the National Institutes of Health.

## Abbreviations and nomenclature

ECL: extracellular loop
MPC: mouse pheochromocytoma
NODAGA: 1,4,7-triazacyclononane-1-glutaric acid-4,7-diacetic acid
pG1: Consomatin pG1
SS-14: somatostatin-14
SS-28: somatostatin-28

