## Supplementary files for "Structure–activity–guided design of potent and selective SSTR2 agonists inspired by a cone snail venom peptide"

**This material includes:**

Fig. S1-10

Table S1-3

```
                N-term                                                      TM1
hSSTR2a         MDMADEPLNGSHTWLSIPFD-----------------LNGSVVSTNTSNQTEPYYDL-TS     42
Dr-sstr2a       ----------MDTWTFMPNSNLSLPDRLVNDSFFPGNESDFGLEFYPPNGTHPGFDH-TS     49
                                                      .         *  .      :.

                                             ICL1 TM2
hSSTR2a         NAVLTFIYFVVCIIGLCGNTLVIYVILRYAKMKTITNIYILNLAIADELFMLGLPFLAMQ     102
Dr-sstr2a       SVVITFVYFVVCAVGLCGNALVIYVILRYAKMKTVTNIYILNLAVADVLCMLSLPFIAIQ     109
                  .:::.:* :** :*****::*****:******* *********:** * **.:**:. .

                   ECL1 TM3                                       ICL2    TM4
hSSTR2a         VALVHWPFGKAICRVVMTVDGINQFTSIFCLTVMSIDRYLAVVHPIKSAKWRRPRTAKMI     162
Dr-sstr2a       LSLLHWPFGSAICRVVLTVDSMNQFTSIFFLTVMSFDRYLAVVHPIKSTKWRKPRMAKTI     169
                  : *****  :*::*:::*.:* ****: * *:*:***::****:*: ::*:*  **

                                     ECL2                TM5
hSSTR2a         TMAVWGVSLLVILPIMIYAGLRSNQWGRSSCTINWPGESGAWYTGFIIYTFILGFLVPLT     222
Dr-sstr2a       SLGMWSVALLVNLPIMIYSGVNAKKNEARTCTMLWPEPQNTYYTAFIFYTFFLGFFLPLI     229
                .:.:* .::** **:::::     :       *.:  *    .  :  : *::*:*::**:.*:

                                ICL3    TM6
hSSTR2a         IICLCYLFIIIKVKSSGIRVGSSKRKKSEKKVTRMVSIVVAVFIFCWLPFYIFNVSSVSM     282
Dr-sstr2a       VISMCYLLIVIKVKSSGMRVGSTKRKRSERKVTRMVSIVVVVFVLCWLPFYVFNVTSVTG     289
                 *.:**::*::*::   .::..   :*:  **:*:*  :*  :**  **::**:**::.::  **

                ECL3 TM7                          H8           C-term
hSSTR2a         AISPTPALKGMFDFVVVLTYANSCANPILYAFLSDNFKKSFQNVLCLVKVSGTDDGERSD     342
Dr-sstr2a       TVPTTPVLKSTFDFVVVLGYANSCANPILYAFLSDNFKKSFQNVLCLKRVGGLDEIDRSD     349
                             :.*:* ***********.***:**::***.:**:        :    .

hSSTR2a         --SK-QDKSRLNETT-----ETQRTLLNGDLQTSI---                           369
Dr-sstr2a       --SR-QDRTRMVNDI--MTETHNAALLNGDLQTTI---                           379
```

**Fig. S1: Sequence alignment of human SSTR2 and Dr-sstr2a.** The N and C-terminus, TM regions, intracellular and extracellular loop (ICL and ECL) regions and Helix 8 (H8) are annotated according to gpcr.db. ECL regions are highlighted in red. Single amino acids that have been reported to interact with SS-14 or octreotide are highlighted in grey. The single amino acid difference within the binding site at TM6 in Dr-sstr2a is highlighted in blue. The conserved DRY and FPFY motifs are highlighted in red.

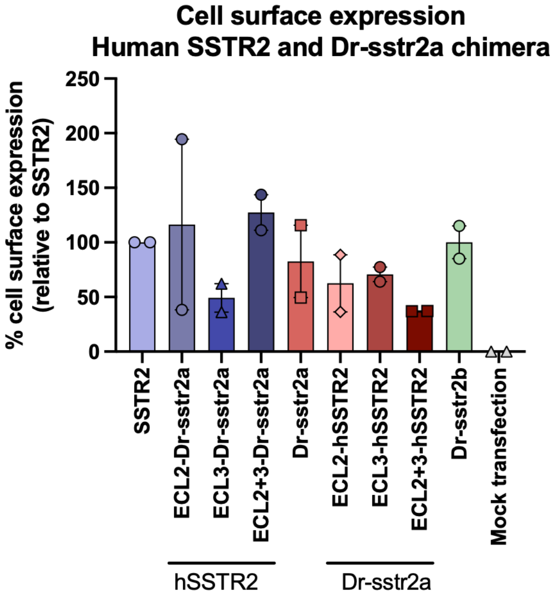


**Fig. S2: Cell surface expression of chimeric human SSTR2 and Dr-sstr2a receptor constructs.** HTLA cells were transiently transfected with different human SSTR2 and Dr-sstr2a chimera, as well as Dr-sstr2b as per the transfection conditions used in the PRESTO-Tango β-arrestin recruitment assay. Means ± S.E.M from two independent biological experiments (each with 8-12 technical replicates) are reported in the bar graph. Mock transfection using empty vector pcDNA3.1+ was done in HTLA cells to determine baseline chemiluminescence signal.

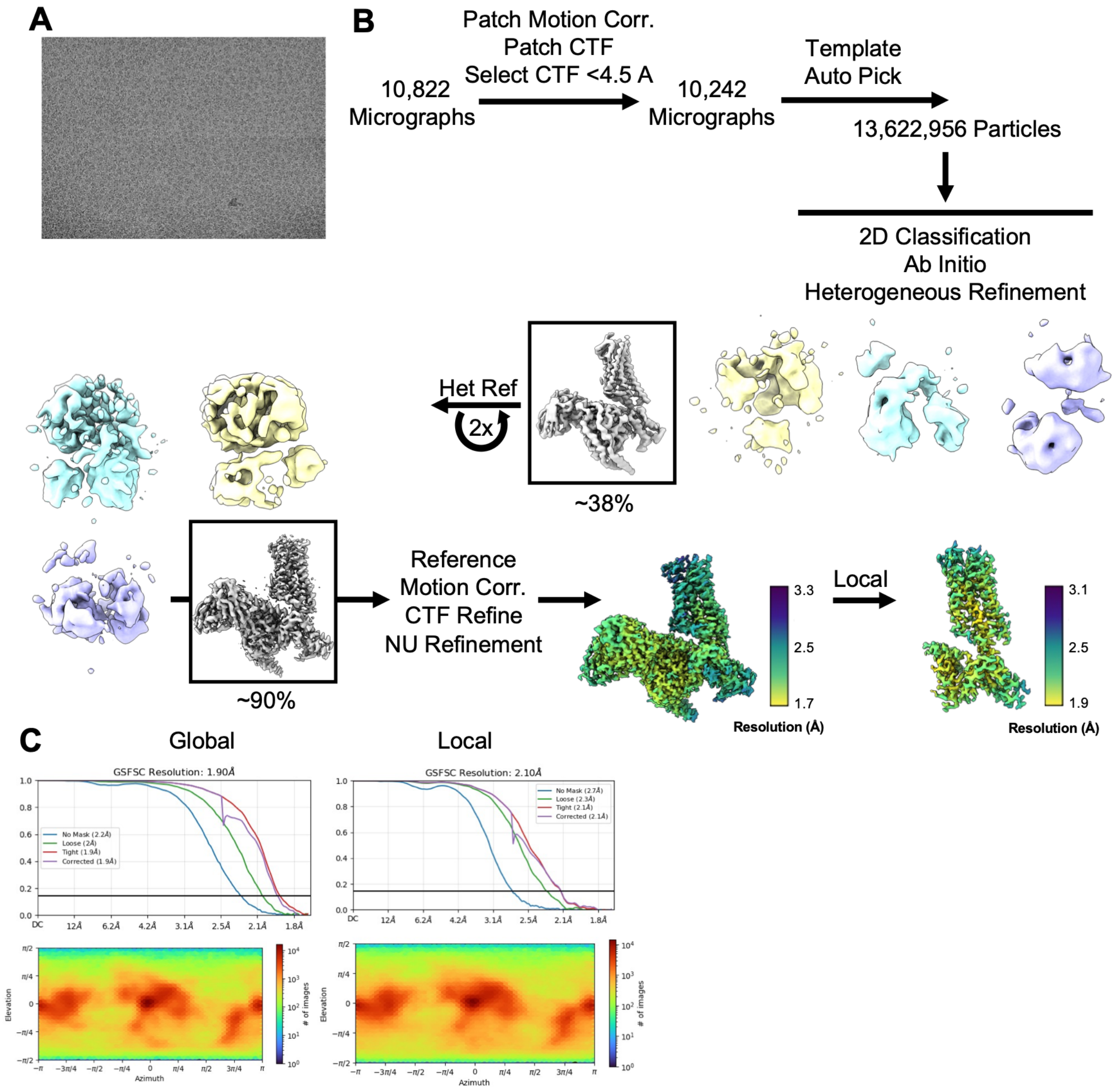


**Fig. S3: SSTR2/Gi3/scFv16/pG1 complex cryo-EM data collection and processing.** (A) Representative cryo-EM micrograph of SSTR2 complex. (B) Cryo-EM processing workflow employed in this study including final maps with local resolution. (C) FSC plots and Euler angle distributions for the maps derived in this work.

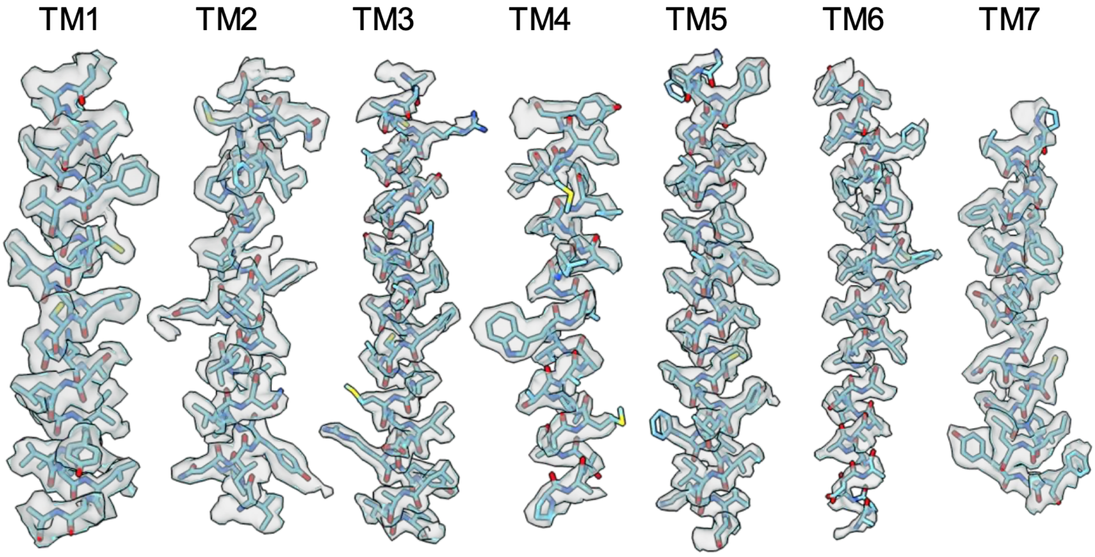


**Fig. S4: Map-Model Agreements.** Map-Model agreement for the SSTR2 transmembrane helices and the local refined receptor map.

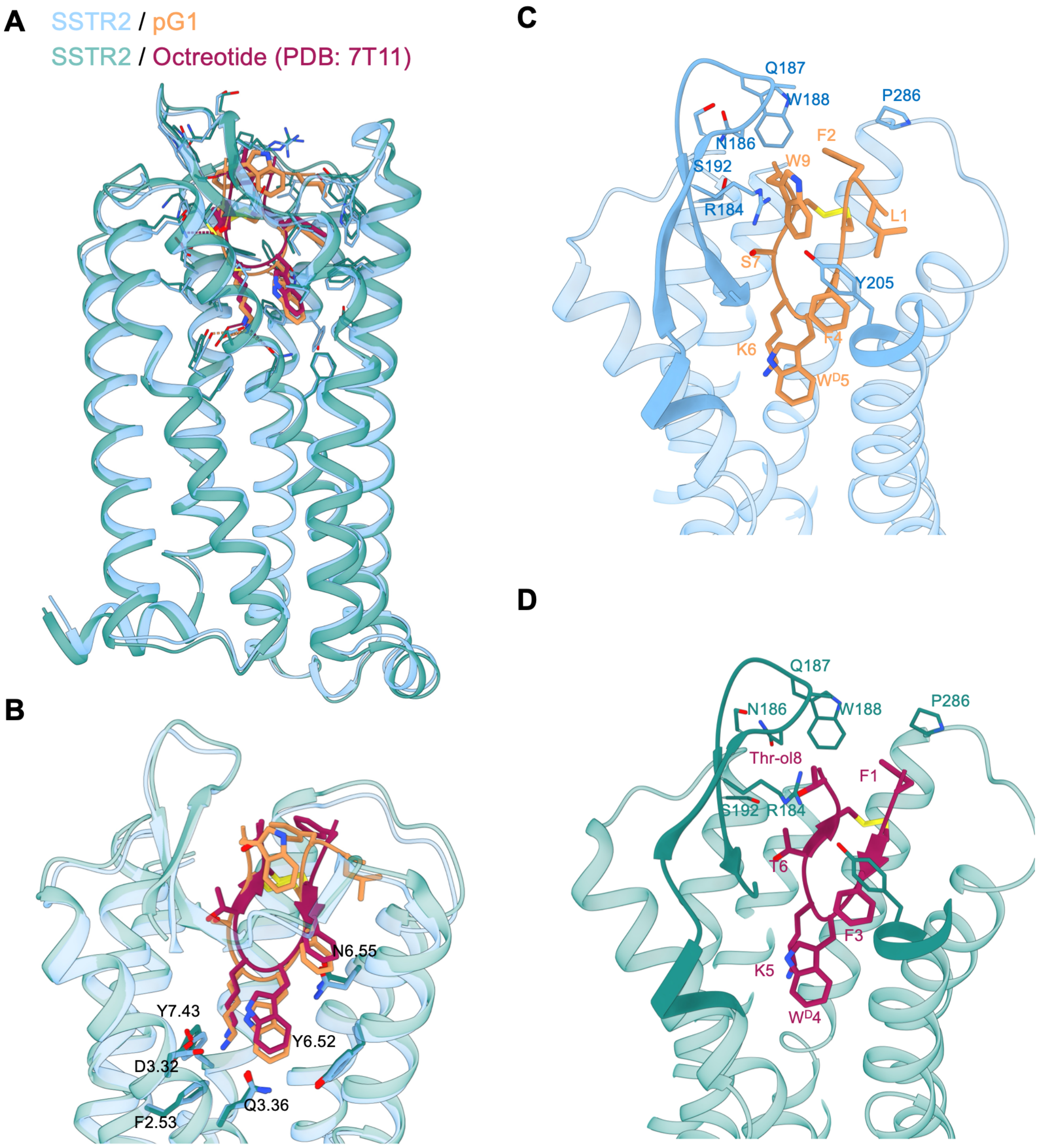


**Fig. S5: Superimposed SSTR2-bound with pG1 or octreotide structures.** (A) Structural overlay of SSTR2/octreotide (pdb: 7T11) and SSTR2/pGi, both in complex with Gi3. (B) Overlaps highlighting the core interactions at the conserved D-Trp and Lys residues of pG1 and octreotide. (C) and (D) show the ECL2 and ECL3 interactions of pG1 and octreotide respectively.

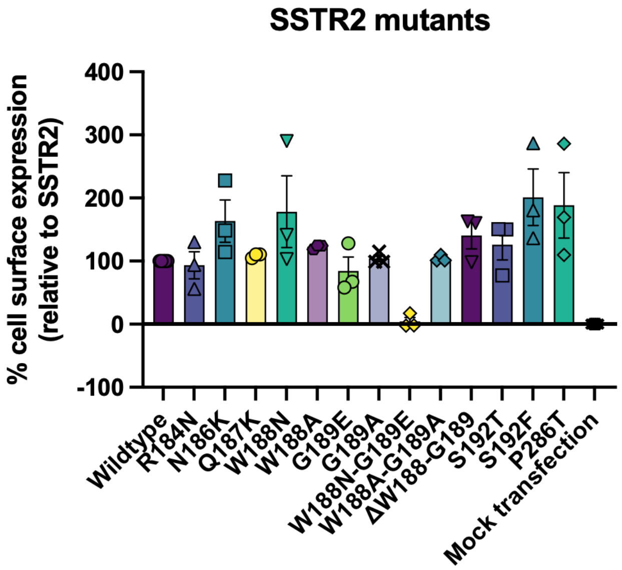


**Fig. S6: Cell surface expression for SSTR2 mutants.** HEK293T cells were transiently transfected with the human wiltype SSTR2 and SSTR2 mutants in the TRUPATH G protein dissociation assay. Means ± S.E.M from three independent experiments (each with 5 technical replicates) are reported in the bar graph. Mock transfection using empty vector pcDNA3.1+ was done in HEK293T cells to determine baseline luminescence signal.

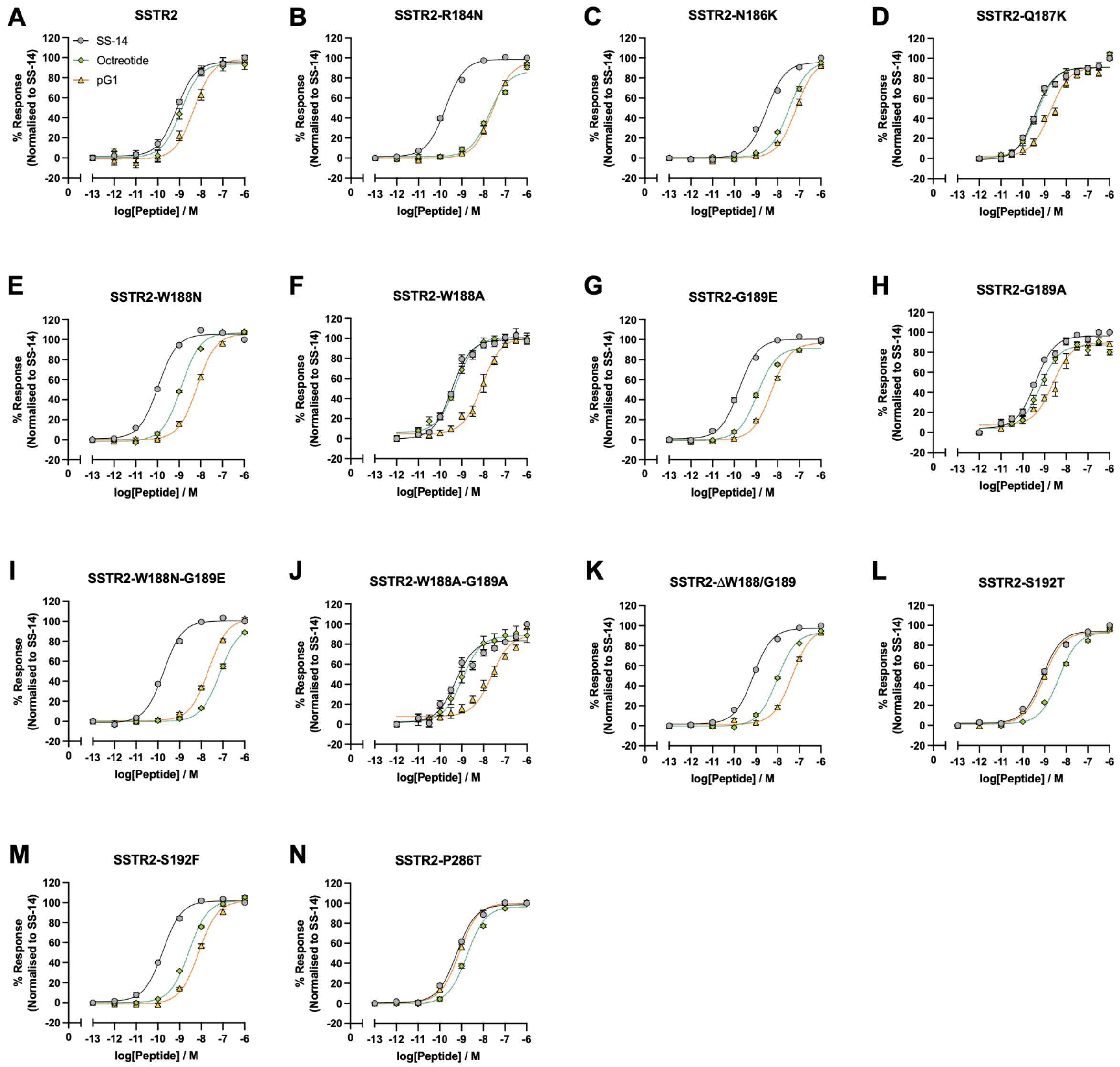


**Fig. S7: SSTR2 ECL2 residues differentially affect pG1 and octreotide G protein dissociation.** (A-N) Concentration-response curves summarizing SSTR2 wild-type and mutants activation by SS-14, octreotide, and pG1 measured in the TRUPATH G protein dissociation assay. HEK293T cells were transiently transfected with SSTR2 wildtype or mutants and Gi3 protein. All data are presented as mean ± S.E.M. from three independent biological replicates, each performed in technical duplicate.

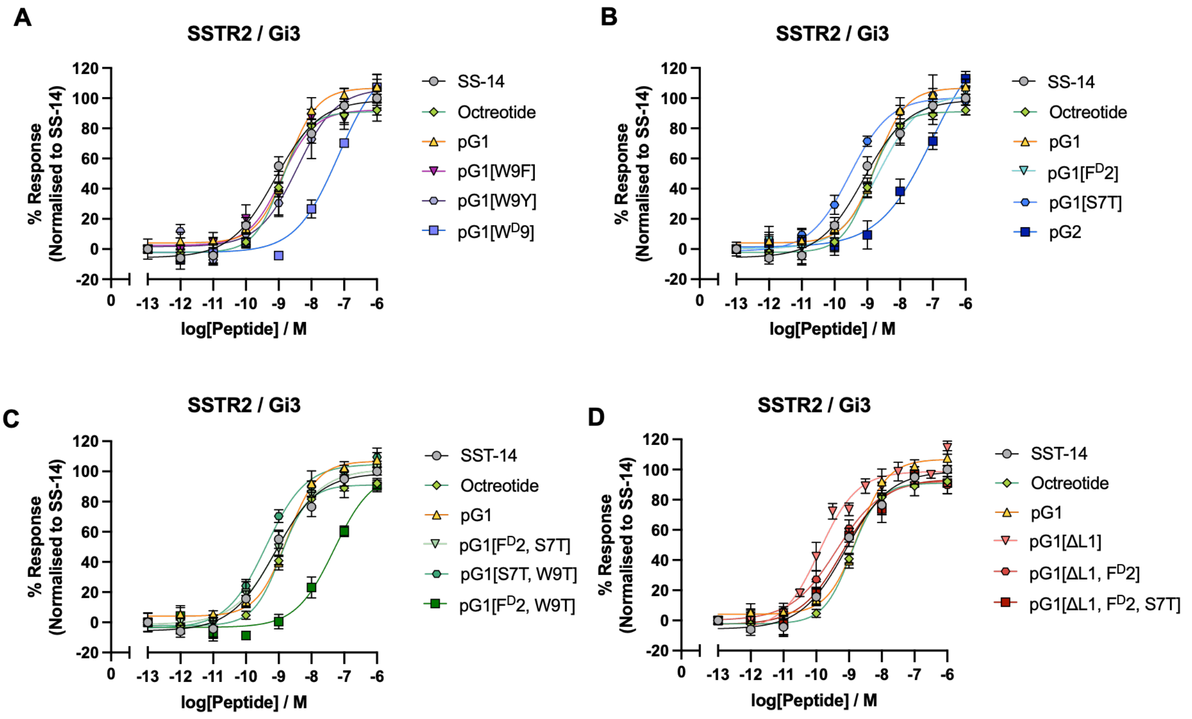


**Fig. S8: pG1 analogs exhibits different extent of SSTR2 activation.** (A-D) Concentration-response curves summarizing SSTR2 activation by a different pG1 analogs measured in the TRUPATH G protein dissociation assay. HEK293T cells were transiently transfected with SSTR2 constructs and Gi3 protein. All data are presented as mean ± S.E.M. from three independent biological replicates, each performed in technical duplicate.

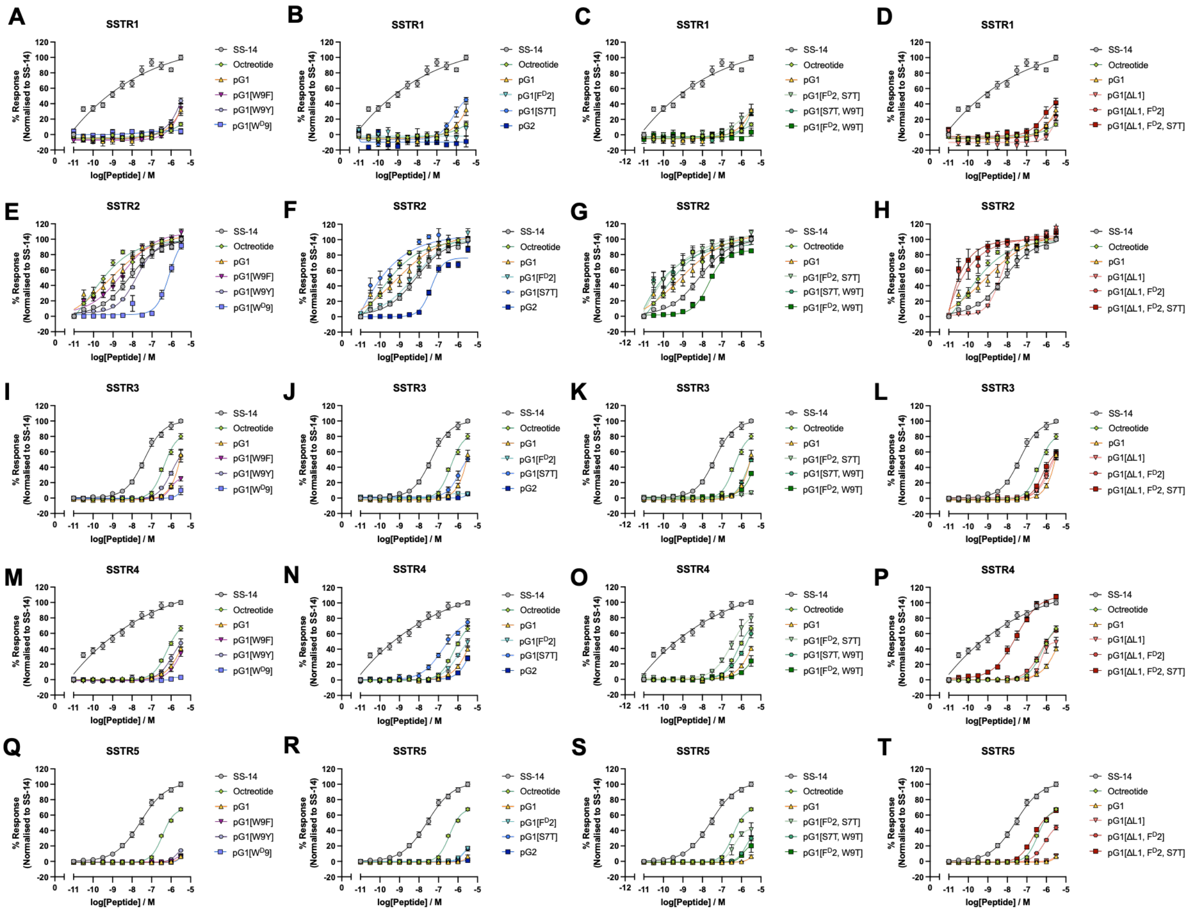


**Fig. S9: pG1 analogs largely retain SSTR2 selectivity.** (A-T) pG1 analogs were tested in the PRESTO-Tango β-arrestin recruitment assay to determine SSTR selectivity. HTLA cells were transiently transfected with human SSTR1-5. All data represented are means ± S.E.M from three independent experiments, each with techical duplicates.

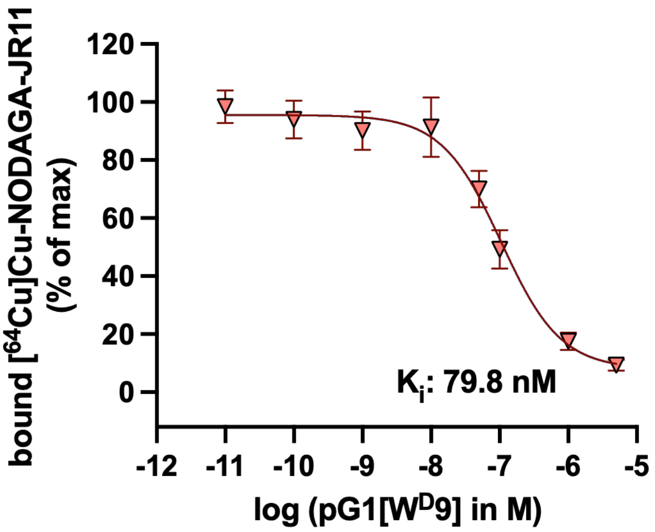


**Fig. S10: pG1[W$^D$9] displays reduced binding affinity at SSTR2.** Representative concentration–response curves showing binding affinities of pG1[W$^D$9] respectively measured using a radiometric competitive binding assay with [$^{64}$Cu]Cu-NODAGA-JR11 as the radiolabeled SSTR2 antagonist in mouse pheochromocytoma cells. All binding data are presented as mean ± S.D. from six independent biological replicates.

**Table S1. Pharmacological parameters of SS-14, octreotide and pG1 mediating SSTRs activation measured by PRESTO-Tango β-arrestin recruitment assay.**

| | Test ligands | **SS-14** | **Octreotide** | **pG1** |
|---|---|---|---|---|
| **$hSSTR_1$** | $pEC_{50}$[a] ± S.E.M. | 9.06 ± 0.60 | N.D. | N.D. |
| | $EC_{50}$ (nM) | 0.879 | | |
| | $E_{max}$[b] ± S.E.M. | 122.91 ± 20.01 | | |
| | Span ± S.E.M. | 168.79 ± 57.03 | | |
| | Hill Slope | 0.21 ± 0.08 | | |
| | n | 11 | 10 | 4 |
| **$hSSTR_2$** | $pEC_{50}$[a] ± S.E.M. | 7.73 ± 0.04 | 9.11 ± 0.05 | 8.95 ± 0.19 |
| | $EC_{50}$ (nM) | 18.47 | 0.780 | 1.14 |
| | $E_{max}$[b] ± S.E.M. | 98.51 ± 1.82 | 95.89 ± 1.22 | 105.66 ± 4.46 |
| | Span ± S.E.M. | 96.41 ± 2.43 | 97.73 ± 2.93 | 114.38 ± 13.00 |
| | Hill Slope | 0.89 ± 0.067 | 0.87 ± 0.07 | 0.40 ± 0.07 |
| | n | 23 | 19 | 12 |
| **$hSSTR_3$** | $pEC_{50}$[a] ± S.E.M. | 6.95 ± 0.09 | 6.31 ± 0.04 | N.D. |
| | $EC_{50}$ (nM) | 112.3 | 493.0 | |
| | $E_{max}$[b] ± S.E.M. | 108.51 ± 5.22 | 83.86 ± 3.35 | |
| | Span ± S.E.M. | 108.19 ± 5.89 | 83.61 ± 3.52 | |
| | Hill Slope | 0.671 ± 0.07 | 1.16 ± 0.46 | |
| | n | 18 | 12 | 8 |
| **$hSSTR_4$** | $pEC_{50}$[a] ± S.E.M. | 8.41 ± 0.43 | N.D. | N.D. |
| | $EC_{50}$ (nM) | 3.93 | | |
| | $E_{max}$[b] ± S.E.M. | 132.13 ± 29.32 | | |
| | Span ± S.E.M. | 164.16 ± 60.99 | | |
| | Hill Slope | 0.21 ± 0.10 | | |
| | n | 10 | 10 | 6 |
| **$hSSTR_5$** | $pEC_{50}$[a] ± S.E.M. | 7.41 ± 0.04 | 6.58 ± 0.05 | N.D. |
| | $EC_{50}$ (nM) | 39.34 | 261.18 | |
| | $E_{max}$[b] ± S.E.M. | 99.48 ± 1.84 | 81.67 ± 3.55 | |
| | Span ± S.E.M. | 99.94 ± 2.27 | 82.04 ± 3.86 | |
| | Hill Slope | 0.78 ± 0.05 | 2.02 ± 5.50 | |
| | n | 19 | 12 | 6 |

Values were generated when the data were fitted to the four-parameter logistic equation. Means ± S.E.M of n individual result sets were shown.

[a] Negative logarithm of agonist concentration when reaching half maximal response.

[b] % of maximal response observed when stimulated with ligands relative to SS-14.

**Table S2. Cryo-EM data collection, refinement and validation statistics.**

| | SSTR2/Gi3/scFv16 Composite, Global (PDB:12LM EMD-76525, 76523) | SSTR2/Gi3/scFv16 Local (EMD-76524) |
|---|---|---|
| **Data collection and processing** | | |
| Voltage (kV) | 300 | 300 |
| Electron exposure (e–/Å2) | 52 | 52 |
| Defocus range (μm) | 0.6 to 1.6 | 0.6 to 1.6 |
| Pixel size (Å) | 0.832 | 0.832 |
| Symmetry imposed | C1 | C1 |
| Initial particle images (no.) | 13,622,956 | 13,622,956 |
| Final particle images (no.) | 2,348,178 | 2,348,178 |
| Map resolution (Å) | 1.9 | 2.1 |
| FSC threshold | 0.143 | 0.143 |
| Map sharpening B factor ($Å^2$) | -58.2 | -67.1 |
| **Refinement** | | |
| Initial Model Used (PDB Code) | 7T11 | |
| Model Resolution | 1.9 | |
| FSC Threshold | 0.5 | |
| *Model Composition* | | |
| Non-hydrogen | 8195 | |
| *B factor ($Å^2$)* | | |
| Protein Atoms | 8195 | |
| Waters & Ions | 0 | |
| *R.M.S Deviations* | | |
| Bonds (Å) | 0.002 | |
| Angles (°) | 0.501 | |
| *Validation* | | |
| MolProbity score | 1.05 | |
| Clashscore | 2.68 | |
| Poor rotamers (%) | 0.92 | |
| *Ramachandran Plot* | | |
| Favored (%) | 98.56 | |
| Allowed (%) | 1.44 | |
| Outliers (%) | 0.00 | |

**Table S3. Pharmacological parameters of SS-14, octreotide, pG1 and its analog in mediating SSTR2 activation measured by TRUPATH G protein dissociation assay.**

| Test ligands | **SS-14** | **Octreotide** | **pG1** |
|---|---|---|---|
| $pEC_{50}$[a] ± S.E.M. | 9.11 ± 0.13 | 8.93 ± 0.13 | 8.75 ± 0.10 |
| $EC_{50}$ (nM) | 0.779 | 1.17 | 1.80 |
| $E_{max}$[b] ± S.E.M. | 98.94 ± 4.48 | 91.03 ± 3.30 | 106.81 ± 4.07 |
| Span ± S.E.M. | 104.67 ± 6.42 | 93.26 ± 4.48 | 102.78 ± 5.26 |
| Hill Slope | 0.64 ± 0.12 | 1.03 ± 0.23 | 0.94 ± 0.18 |
| n | 8 | 6 | 8 |

| Test ligands | **pG1[W9F]** | **pG1[W9Y]** | **pG1[$W^D$9]** |
|---|---|---|---|
| $pEC_{50}$[a] ± S.E.M. | 8.91 ± 0.20 | 8.12 ± 0.27 | 7.19 ± 0.25 |
| $EC_{50}$ (nM) | 1.24 | 7.68 | 65.3 |
| $E_{max}$[b] ± S.E.M. | 92.25 ± 6.64 | 100.71 ± 11.18 | 124.99 ± 17.93 |
| Span ± S.E.M. | 89.86 ± 8.97 | 102.16 ± 13.4 | 127.34 ± 18.89 |
| Hill Slope | 0.87 ± 0.33 | 0.61 ± 0.21 | 0.67 ± 0.18 |
| n | 6 | 6 | 4 |

| Test ligands | **pG1[$F^D$2]** | **pG1[S7T]** | **pG2** |
|---|---|---|---|
| $pEC_{50}$[a] ± S.E.M. | 8.63 ± 0.20 | 9.53 ± 0.17 | 7.02 ± 0.46 |
| $EC_{50}$ (nM) | 2.33 | 0.30 | 95.2 |
| $E_{max}$[b] ± S.E.M. | 102.48 ± 7.54 | 100.41 ± 4.89 | 144.00 ± 32.91 |
| Span ± S.E.M. | 102.18 ± 9.68 | 101.87 ± 7.64 | 142.72 ± 34.58 |
| Hill Slope | 0.70 ± 0.20 | 0.71 ± 0.17 | 0.53 ± 0.16 |
| n | 7 | 6 | 6 |

| Test ligands | **pG1[$F^D$2, S7T]** | **pG1[S7T, W9T]** | **pG1[$F^D$2, W9T]** |
|---|---|---|---|
| $pEC_{50}$[a] ± S.E.M. | 9.01 ± 0.18 | 9.39 ± 0.11 | 7.31 ± 0.20 |
| $EC_{50}$ (nM) | 0.98 | 0.41 | 48.8 |
| $E_{max}$[b] ± S.E.M. | 101.48 ± 6.02 | 104.83 ± 3.48 | 99.70 ± 11.57 |
| Span ± S.E.M. | 102.88 ± 8.42 | 108.74 ± 5.25 | 103.07 ± 12.46 |
| Hill Slope | 0.65 ± 0.16 | 0.72 ± 0.11 | 0.74 ± 0.19 |
| n | 8 | 7 | 6 |

| Test ligands | **pG1[ΔL1]** | **pG1[ΔL1, $F^D$2]** | **pG1[ΔL1, $F^D$2, S7T]** |
|---|---|---|---|
| $pEC_{50}$[a] ± S.E.M. | 9.78 ± 0.28 | 9.38 ± 0.17 | 9.20 ± 0.17 |
| $EC_{50}$ (nM) | 0.17 | 0.41 | 0.63 |
| $E_{max}$[b] ± S.E.M. | 105.21 ± 6.73 | 92.87 ± 4.58 | 93.56 ± 5.26 |
| Span ± S.E.M. | 117.85 ± 17.84 | 92.85 ± 7.05 | 96.27 ± 7.67 |
| Hill Slope | 0.50 ± 0.16 | 0.63 ± 0.14 | 0.67 ± 0.17 |
| n | 6 | 6 | 6 |

Values were generated when the data were fitted to the four-parameter logistic equation. Means ± S.E.M of n individual result sets were shown.

[a] Negative logarithm of agonist concentration when reaching half maximal response.

[b] % of maximal response observed when stimulated with ligands relative to SS-14.

### HPLC and MS spectra for synthesized pG1 analogues

### pG1[W9F]

LF**C**FwKS**C**F

Purified by preparative RP-HPLC, 25-50% A:B over 30 min. 11 mg, 28%, 90.8% purity. HRMS (ESI) (m/z): $[M+H]^+$ calcd. for $C_{59}H_{76}N_{11}O_{11}S_2$ 1179.4473; found 1179.5164.

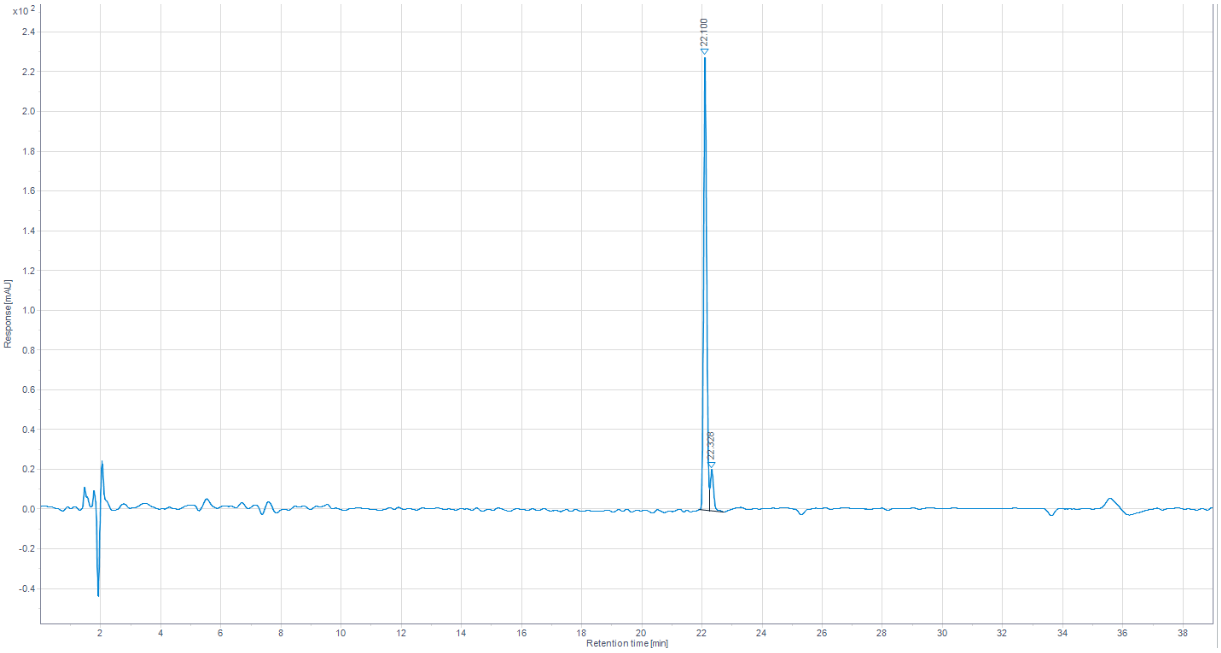


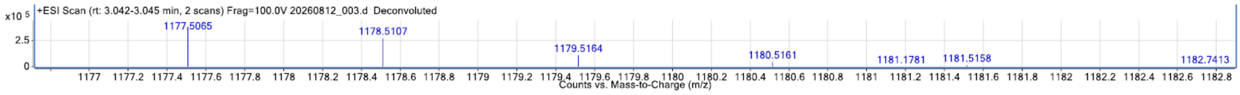

**pG1[W9Y]**

LF**C**FwKS**C**Y

Purified by preparative RP-HPLC, 25-50% A:B over 30 min. 8 mg, 23%, 90.1% purity. HRMS (ESI) (m/z): $[M+H]^+$ calcd. for $C_{59}H_{76}N_{11}O_{12}S_2$ 1195.4373; found 1195.5123.

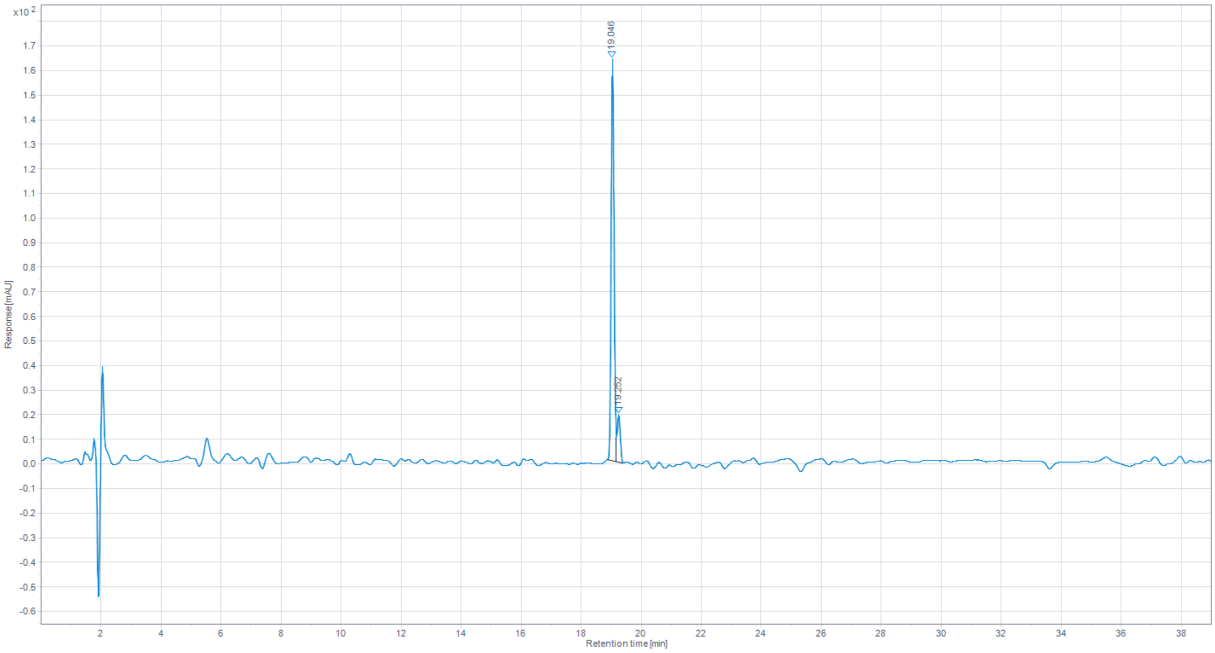


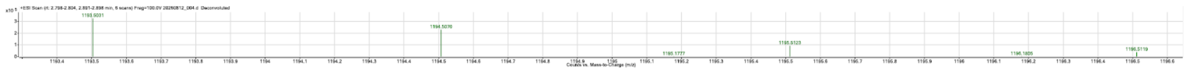

**pG1[W$^{D}$9]**

LF**C**FwKS**C**w

Purified by preparative RP-HPLC, 25-50% A:B over 30 min. 2 mg, 7%, >99.9% purity. HRMS (ESI) (m/z): $[M+H]^+$ calcd. for $C_{61}H_{77}N_{12}O_{11}S_2$ 1217.5265; found 1217.5268.

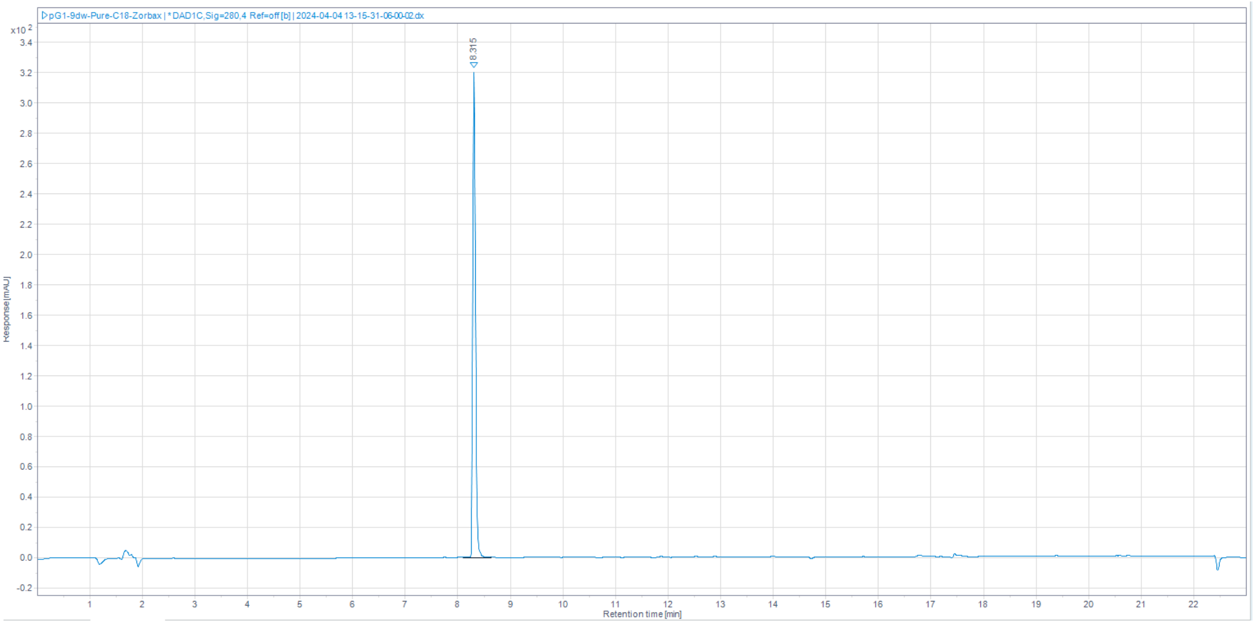


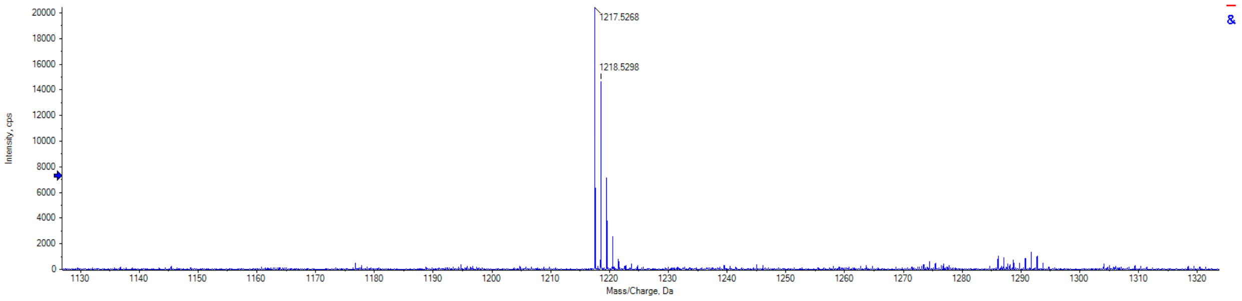

**pG1[F$^D$2]**

LfCFwKSCW

Purified by preparative RP-HPLC, 25-50% A:B over 25 min. 23 mg, 41%, 96.6% purity. HRMS (ESI) (m/z): $[M+H]^+$ calcd. for $C_{61}H_{77}N_{12}O_{11}S_2$ 1217.5271; found 1217.5269.

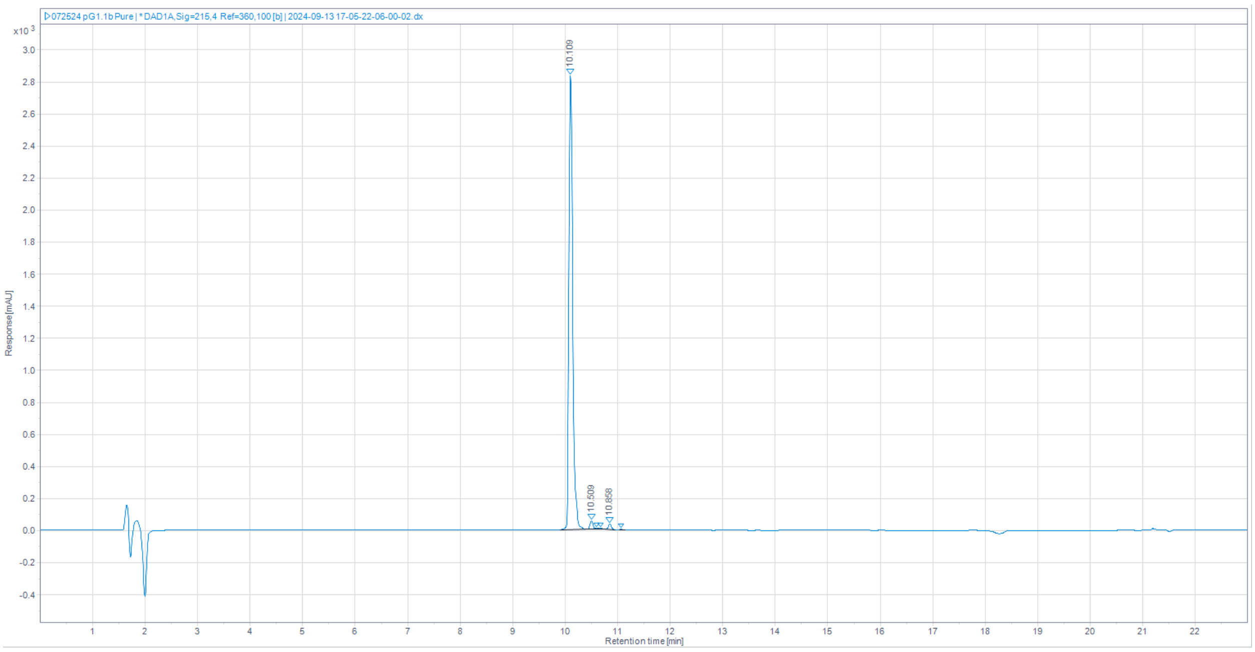


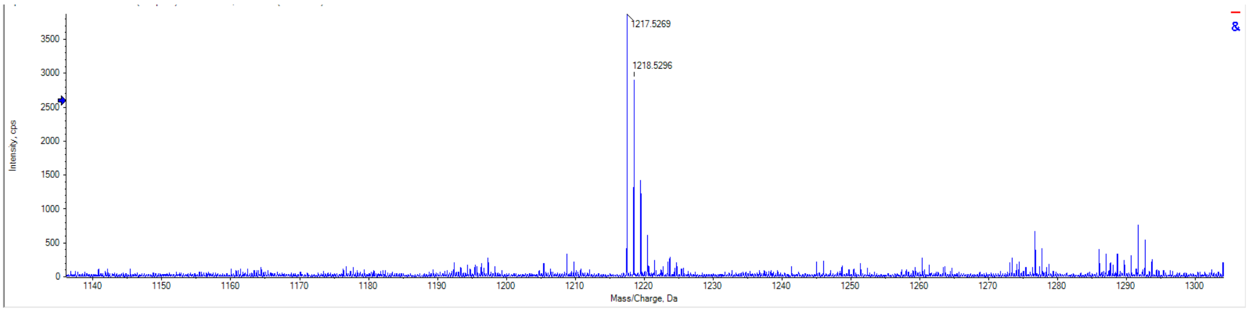

**pG1[S7T]**

LF**C**FwKS**C**W

Purified by preparative RP-HPLC, 25-50% A:B over 25 min. 6 mg, 16%, >99.9% purity. HRMS (ESI) (m/z): $[M+H]^+$ calcd. for $C_{61}H_{77}N_{12}O_{11}S_2$ 1217.5271; found 1217.5269.

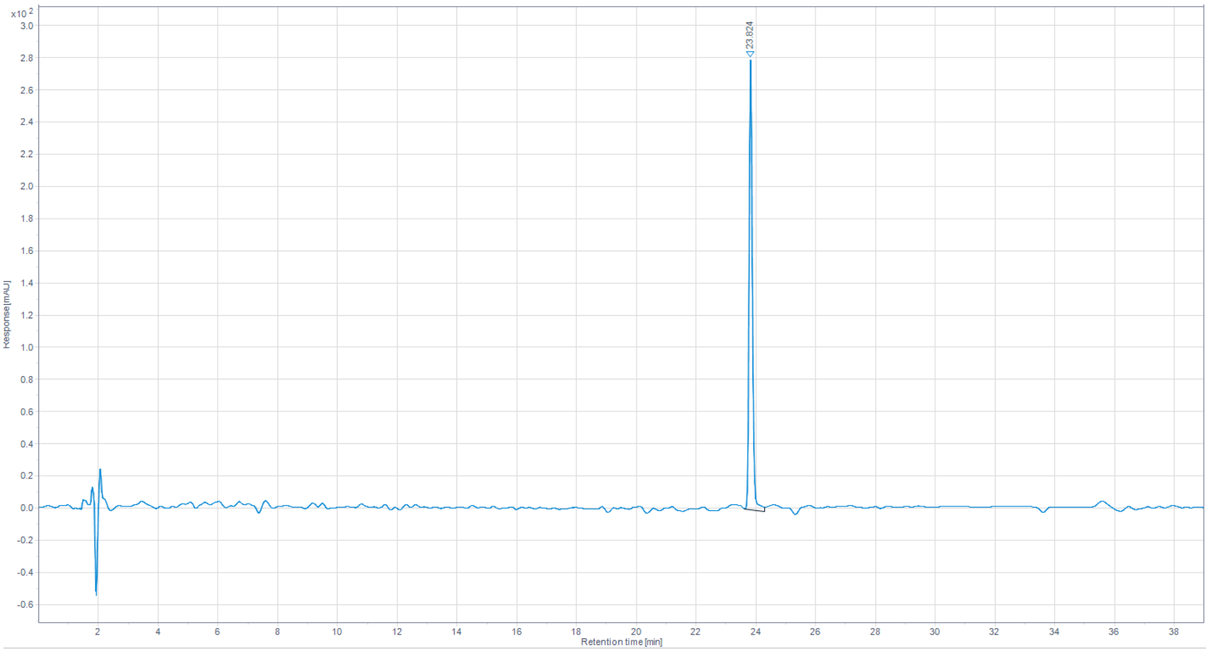


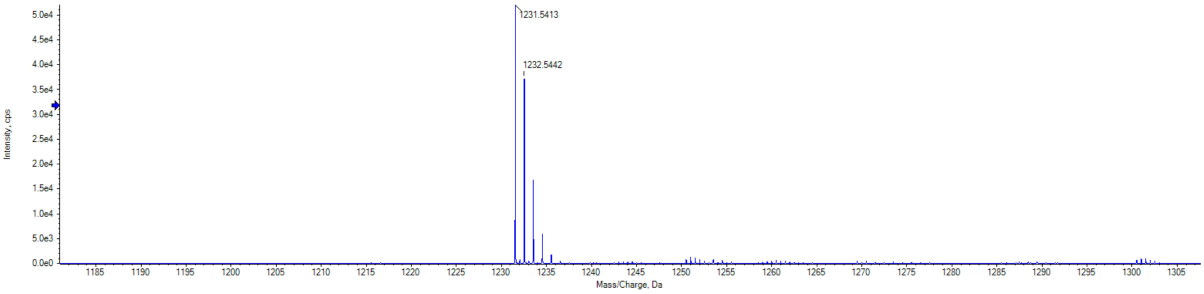

**pG1[F$^D$2, S7T]**

LfCFwKTCW

Purified by preparative RP-HPLC, 25-50% A:B over 30 min. 4 mg, 11%, 99.9% purity. HRMS (ESI) (m/z): $[M+H]^+$ calcd. for $C_{62}H_{79}N_{12}O_{11}S_2$ 1231.5422; found 1231.5413.

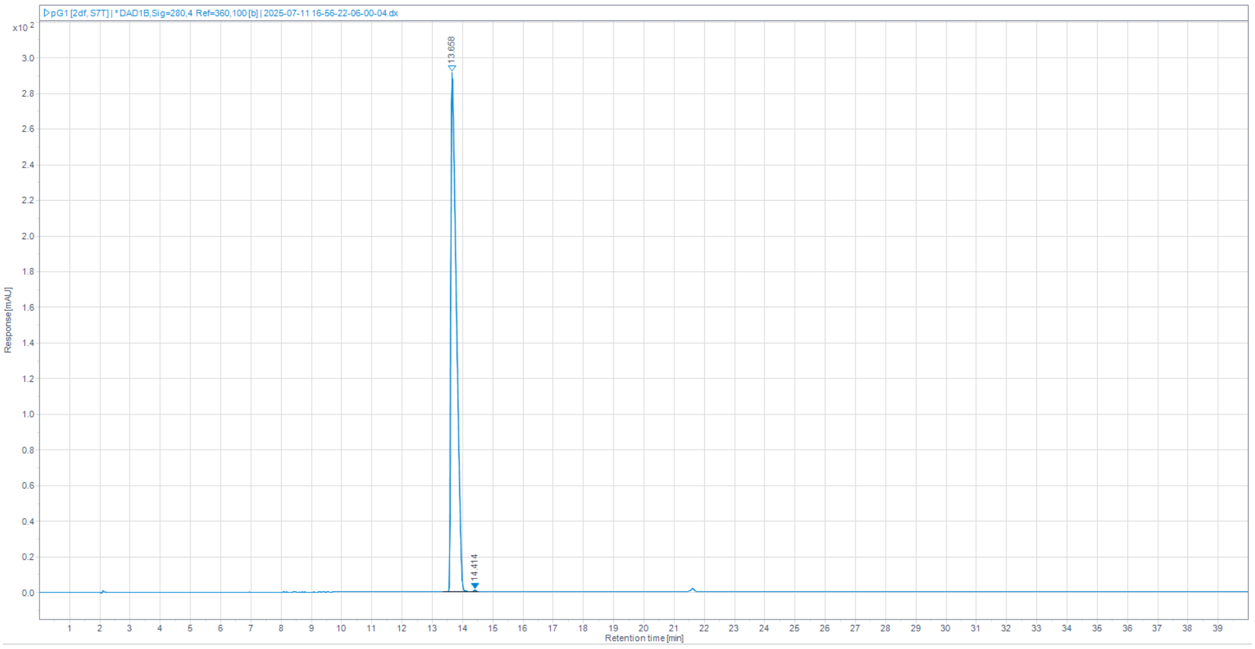


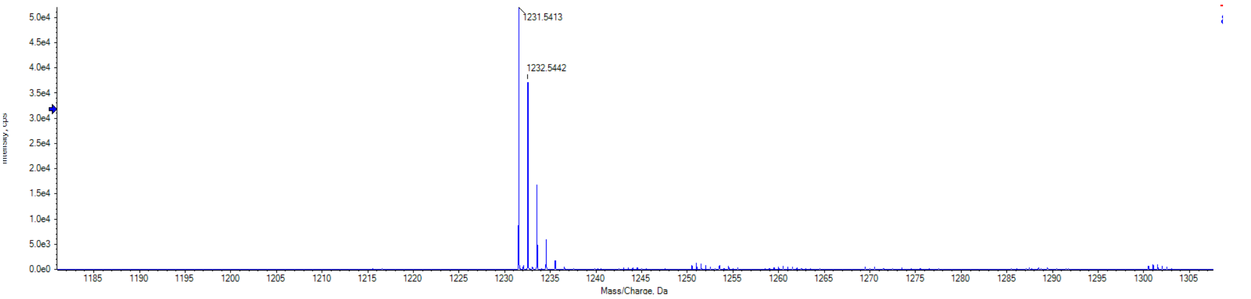

**pG1[S7T, W9T]**

LF**C**FwKT**C**T

Purified by preparative RP-HPLC, 25-50% A:B over 30 min. 5 mg, 15%, 95.1% purity. HRMS (ESI) (m/z): $[M+H]^+$ calcd. for $C_{55}H_{76}N_{11}O_{12}S_2$ 1146.5105; found 1146.5095.

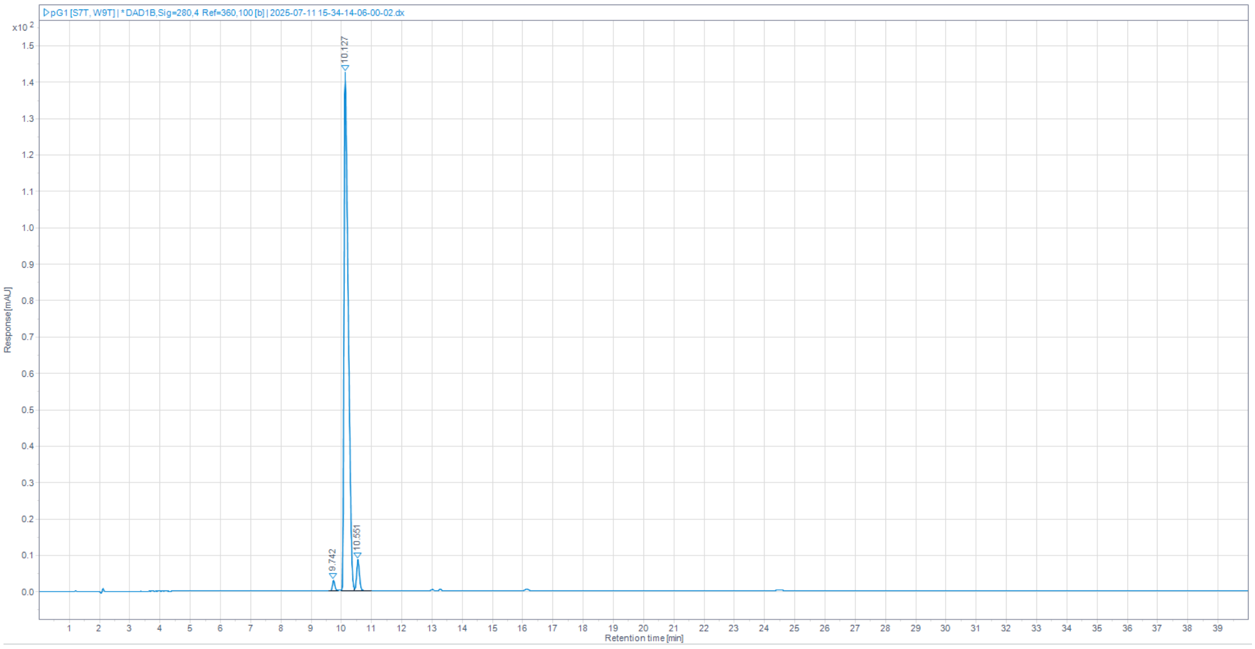


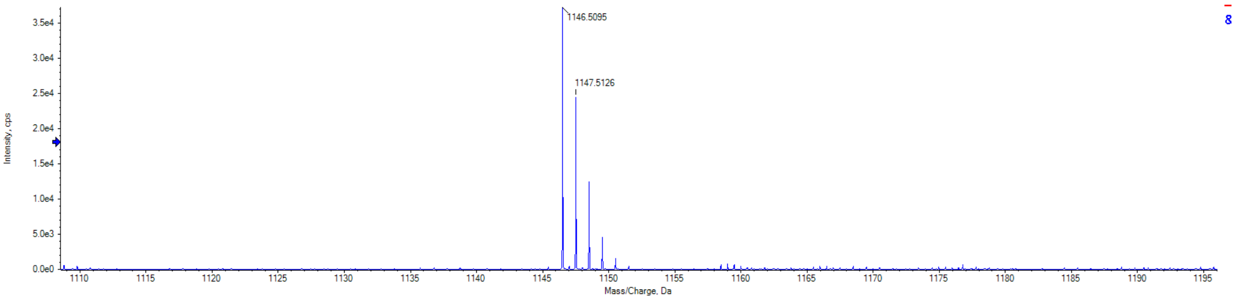

**pG1[F^D^2, W9T]**

LfCFwKSCT

Purified by preparative RP-HPLC, 33-40% A:B over 30 min. 2 mg, 6%, 99.7% purity. HRMS (ESI) (m/z): $[M+H]^+$ calcd. for $C_{54}H_{74}N_{11}O_{12}S_2$ 1132.4949; found 1132.4948

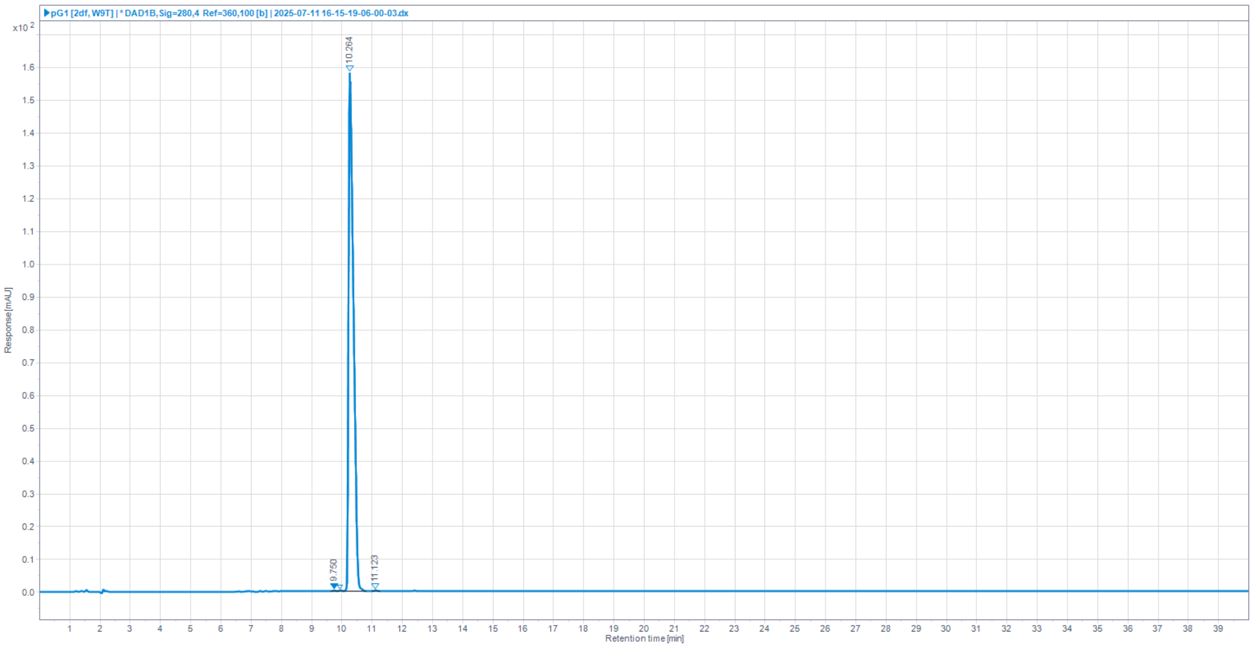


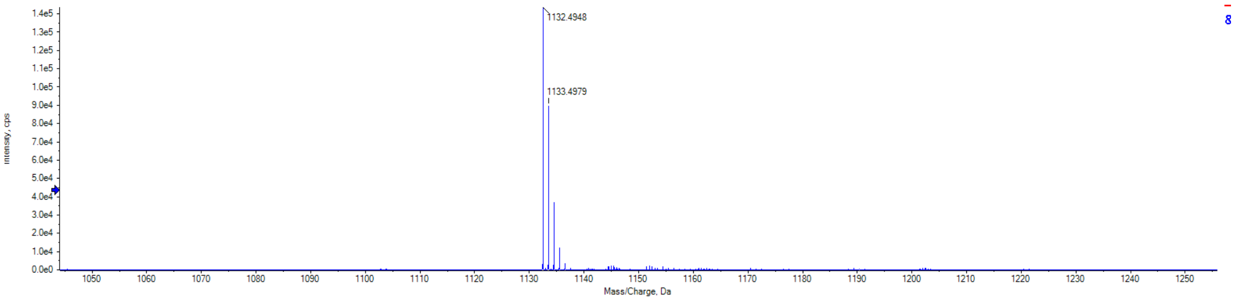

**pG1[∆L1]**

FCFwKSCW

Purified by preparative RP-HPLC, 30-45% A:B over 25 min. 5 mg, 16%, 98.9% purity. HRMS (ESI) (m/z): $[M+H]^+$ calcd. for $C_{55}H_{66}N_{11}O_{10}S_2$ 1104.4425; found 1104.4433.

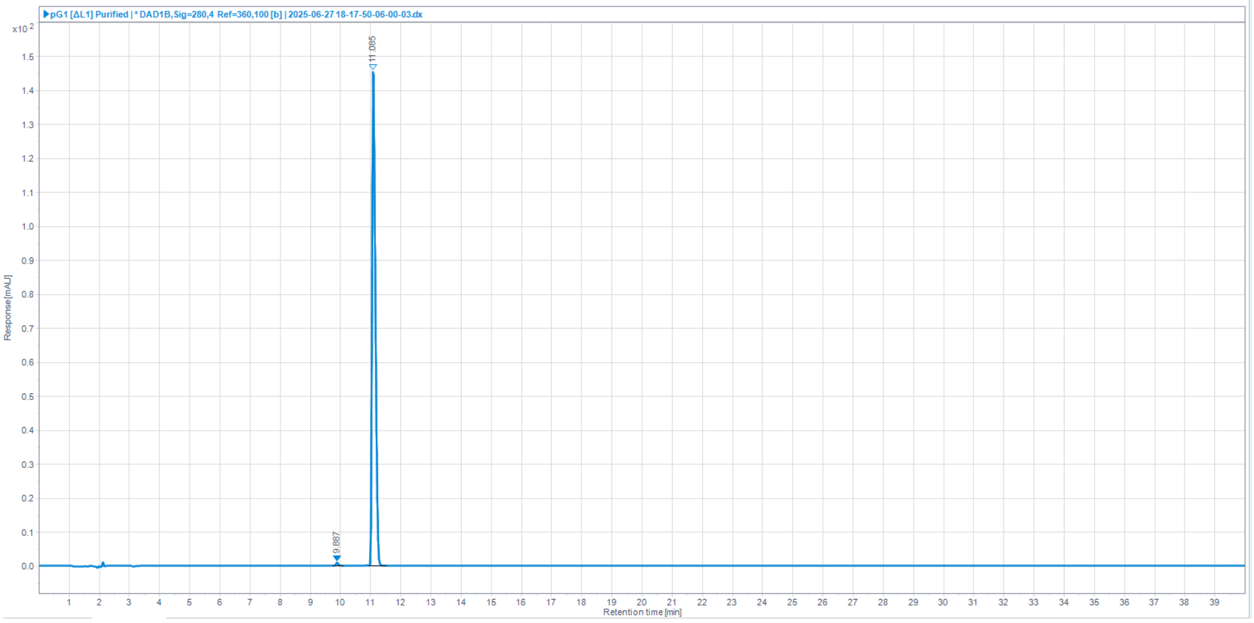


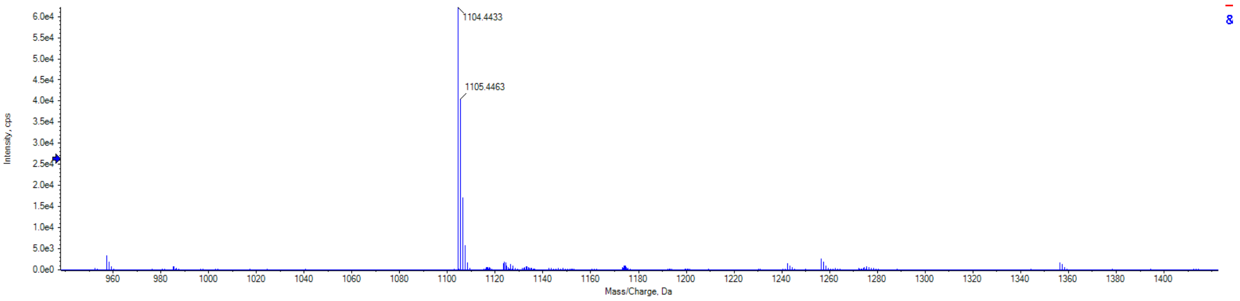

**pG1[ΔL1, F^D^2]**

fCFwKSCW

Purified by preparative RP-HPLC, 37-42% A:B over 30 min. 0.1 mg, 0.4%, 99.8% purity. HRMS (ESI) (m/z): $[M+H]^+$ calcd. for $C_{55}H_{66}N_{11}O_{10}S_2$ 1104.4425; found 1104.4437.

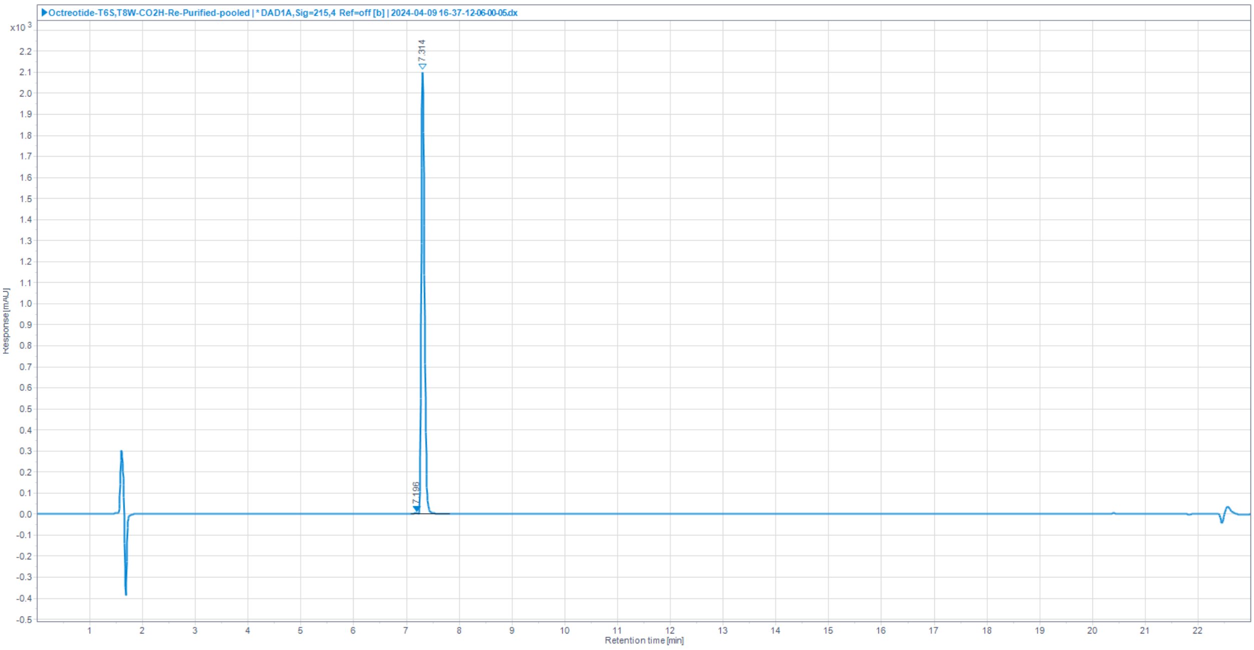


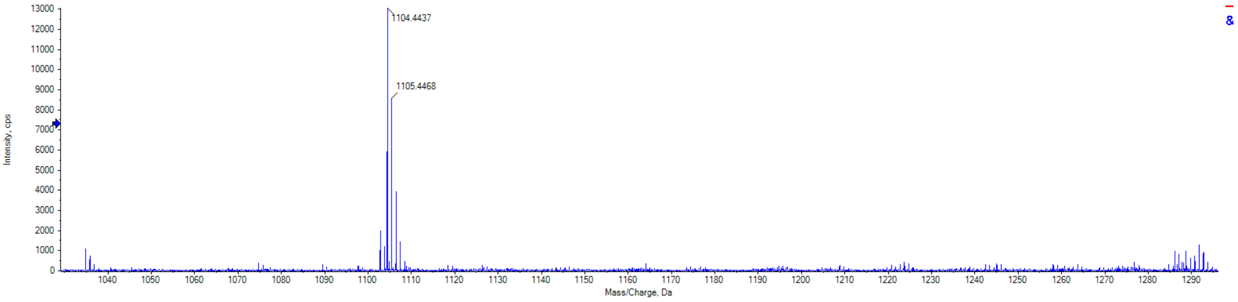

**pG1[ΔL1, F$^{D}$2, S7T]**

f**C**FwKT**C**W

Purified by preparative RP-HPLC, 39-46% A:B over 30 min. 1 mg, 4%, 98.3% purity. HRMS (ESI) (m/z): [M+H]$^{+}$ calcd. for $C_{56}H_{68}N_{11}O_{10}S_2$ 1118.4581; found 1118.4577.

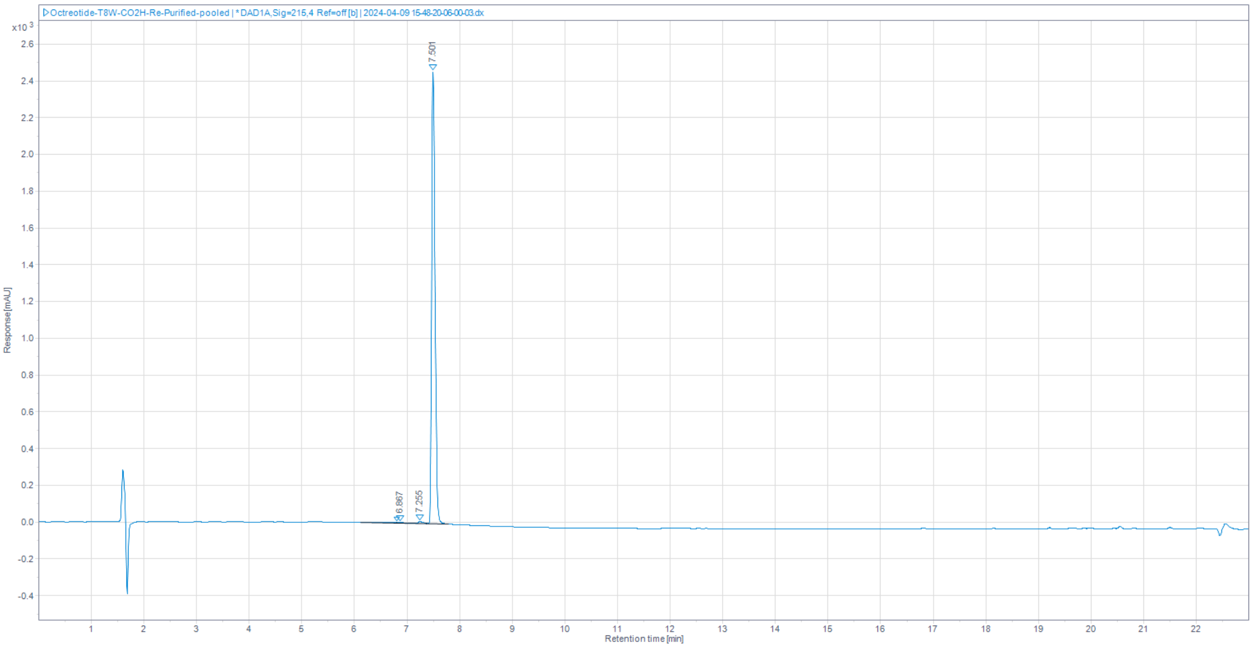


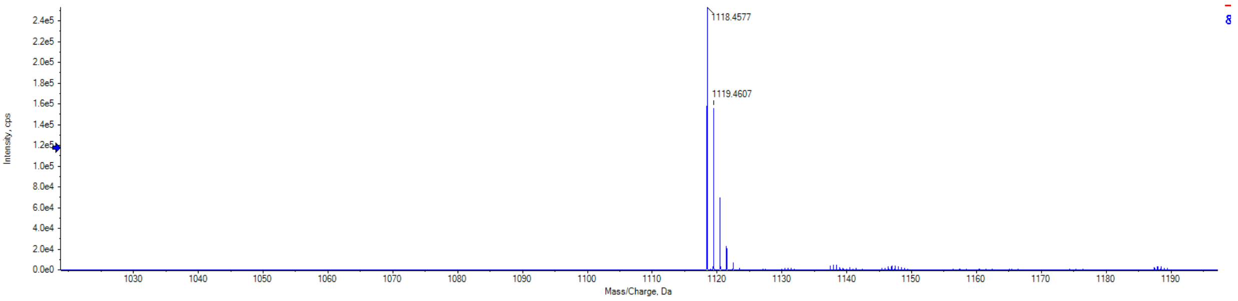
